# Spatiotemporal photocatalytic proximity labeling proteomics reveals ligand-activated extracellular and intracellular EGFR neighborhoods

**DOI:** 10.1101/2024.12.21.629933

**Authors:** Zhi Lin, Wayne Ngo, Yu-Ting Chou, Harry Wu, Katherine J. Susa, Young-wook Jun, Trever G. Bivona, Jennifer A. Doudna, James A. Wells

## Abstract

Photo-proximity labeling proteomics (PLP) methods have recently shown that the cell surface receptors can dynamically form lateral interactome networks. Here, we present a paired set of PLP workflows that simultaneously track neighborhood changes for oncogenic epidermal growth factor receptor (EGFR) with temporal resolution, both outside and inside of cells. We achieved this by augmenting the multiscale PLP workflow we call MultiMap, where three photo-probes with different labeling ranges were photo-activated by one photocatalyst, Eosin Y. By anchoring Eosin Y extracellularly and intracellularly on EGFR, we captured hundreds of proteins on both sides of the cell membrane that change in proximity to EGFR upon EGF activation. Neighbors engaged with EGFR within minutes to over an hour, reflecting dynamic interactomes during early, mid- and late-signaling including phosphorylation, internalization, degradation and transcriptional regulation. This rapid “photographic” labeling approach provides snapshots of signaling neighborhoods, revealing their dynamic nature, and potential for drug targeting.

## Introduction

Epidermal growth factor receptor (EGFR) is a classic paradigm for growth factor signaling.^1^ Seminal studies over the past decade have established EGFR as one of the first transmembrane receptors to be mechanistically and structurally characterized to show that ligand-receptor recognition is critically tied to multiple cytosolic signaling steps with different time-scales.^2-7^ EGFR is also a key therapeutic node centric in cancer signaling. Rampant dysregulation of EGFR is frequently observed in various cancer malignancies, including non-small cell lung cancer (NSCLC) and breast cancer.^8, 9^ Antibody and small molecule therapeutics targeting the extracellular domain or kinase domain of EGFR have proven effective in delaying cancer progression.^10-12^ However, cancers invariably develop resistance to these receptor-oriented treatments and the presence of EGFR on normal tissues can create dose-limiting toxicity.^13, 14^ Thus, targeting functional EGFR interactions may provide an alternative strategy.

Proteins in the cell membrane are densely packed at distances estimated no more than 60-70Å apart, which is about the girth of an average protein.^15-17^ The 2D environment in the membrane allows weak complexes to form rapidly and stably even when intrinsic dissociation constants can be 2-3 logs lower than 3D counterparts.^18^ These physical considerations have led to the development of photo-proximity labeling proteomics (PLP) using highly reactive carbene intermediates such as μMap.^19^ In this method, an iridium photocatalyst conjugated to an antibody to trigger an aryl-diazirine-biotin photo-probe to form carbene species with an estimated labeling radius of ∼110Å from a single conjugation site on the primary antibody.^20^ We recently reported the development of MultiMap,^21^ a multiscale photocatalytic PLP platform that utilizes a single commercially available photocatalyst, Eosin Y, to trigger labeling at multiple ranges from ∼100-3000Å using three different photo-probes containing either an aryl-diazirine, aryl-azide or phenol warhead.^22^ Using MultiMap, we conjugated Eosin Y to cetuximab, a clinically approved EGFR antibody that competes with its natural ligand EGF and blocks EGFR signaling,^23, 24^ and probed this off-state neighborhood of EGFR with adjustable resolution. However, in order to map functional EGFR neighborhoods, we sought a universal way to label surface proteins with minimal impact on ligand stimulation events from both inside and outside the cells to allow for the study of dynamic interactions and signaling processes at multiple resolutions.

Here, we introduce a dual PLP approach where one can interrogate membrane-associated neighborhoods at multiple resolutions from the extracellular or intracellular side of the receptor that we call eMultiMap and iMultiMap, respectively. We engineered the full-length EGFR into cells fused with an extracellular N-terminal Flag-Tag and an intracellular C-terminal HaloTag without affecting signaling. The EY photocatalyst was readily anchored using an EY-conjugated anti-Flag antibody for eMultiMap, or EY-HaloTag Ligand for iMultiMap. By adding EGF and triggering PLP reactions with blue light at different timepoints, we identified more than 300 proteins whose labeling patterns changed over a period of 5 min to 1 hour. We validated ten candidates most highly associated with EGFR activation that included phosphatases, trafficking proteins and transcription factors, reflecting early, mid- and late-signaling processes. This approach is somewhat analogous to molecular photography which captures transient EGFR signaling complexes that may inspire targets for drug discovery.

## Results

### Genetically encoded tags allow selective labeling in live cells in a spatiotemporally controlled manner

To label EGFR on either side of the membrane with the EY photocatalyst (**Figure 1A**), we fused a Flag-tag on the N-terminus of the extracellular domain (ECD) of EGFR and a HaloTag on the C-terminus of the intracellular domain (ICD), designated Flag-EGFR and EGFR-HaloTag (Flag-EGFR-HaloTag), respectively (**Figure S1A**). We introduced these into A549 cells, a non-small cell lung cancer cell line, for its well-established and robust response to EGF activation as well as its physiologically relevant expression level of EGFR.^25, 26^ After transfecting the two constructs in A549 cells, we saw significant shifts for the anti-Flag signal on cells for both constructs by flow cytometry (**Figure S1B**). We adjusted transfection conditions so that both engineered EGFR constructs, Flag-EGFR and EGFR-HaloTag, are at comparable levels to the endogenous EGFR (**Figure 1B**).

**Figure 1:**
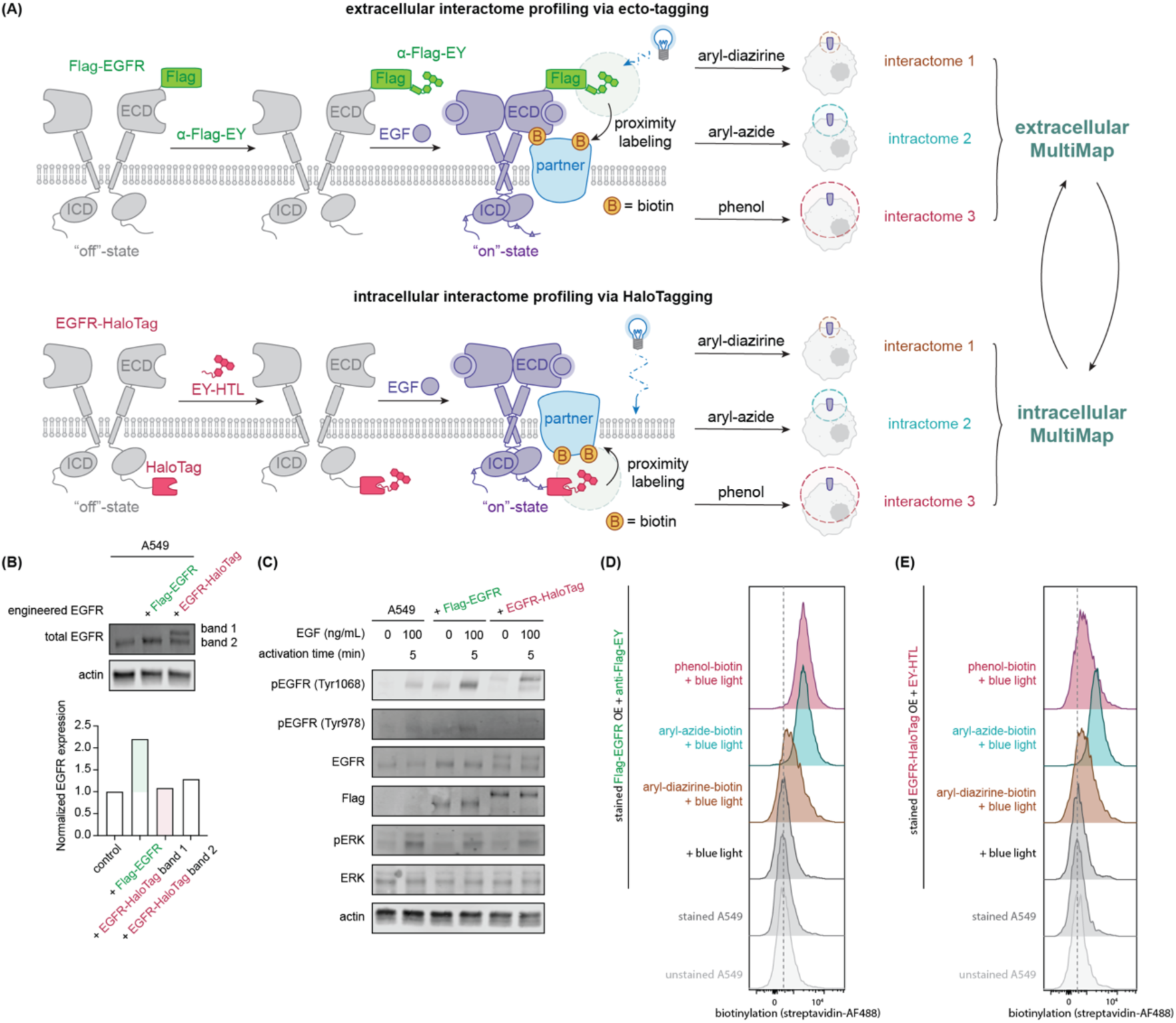
Systematic interactome profiling in live cells enabled by extracellular MultiMap (eMultiMap) and intracellular MultiMap (iMultiMap). (A) Schematics of eMultiMap and iMultiMap workflows enabling photocatalytic biotinylation of neighboring proteins from either the extracellular domain (ECD) or intracellular domain (ICD) of EGFR, respectively. The photocatalyst, Eosin Y (EY), is introduced to EGFR by incubating cells expressing an N-terminal Flag-tag (Flag-EGFR) or a C-terminal HaloTag (EGFR-HaloTag); EY is anchored using an EY-conjugated anti-Flag antibody or a derivatized EY with a HaloTag Ligand, respectively. Cells are illuminated for 2 min with blue light in the presence of three different biotinylated photo-probes of increasing labeling distances (aryl-diazirine-biotin, aryl-azide-biotin and phenol-biotin). EGFR and its neighbors are biotinylated, enriched and digested for MS analysis. These photo-probes allowed for tracking dynamic neighborhoods under different conditions, including EGF-triggered activation. **(B)** For eMultiMap and iMultiMap workflows, Flag-EGFR and EGFR-HaloTag were expressed in the cells at roughly equimolar amounts relative to endogenous EGFR as detected by western blot (WB) analysis. The small size difference between the Flag-EGFR and endogenous EGFR cannot be resolved by WB; EGFR-HaloTag is 33 kDa larger compared to endogenous EGFR, resulting in an upshifted band 1. **(C)** Engineered EGFR constructs respond indistinguishably to EGF stimuli (100 ng/μL) as seen in EGFR phosphorylation and downstream ERK phosphorylation. **(D)** On-cell labeling of Flag-EGFR with aryl-diazirine-biotin, aryl-azide-biotin and phenol-biotin in the presence of EY-conjugated Flag antibody (anti-Flag-EY) and blue light illumination. **(E)** On-cell labeling of EGFR-HaloTag with aryl-diazirine-biotin, aryl-azide-biotin and phenol-biotin in the presence of EY-HaloTag Ligand (EY-HTL) and blue light illumination.

In order to validate that the tagged EGFR constructs functioned properly upon EGF ligand activation, we incubated the engineered cells with EGF and observed significant activation of well-known and functional phosphorylation sites on EGFR, Tyr1086 and Tyr978,^27^ at comparable levels to the wild-type cells (**Figure 1C**).We also confirmed that the cell engineering did not significantly alter the patterns of downstream ERK levels or ERK phosphorylation upon EGF activation.

We then showed that both the Flag-tag and HaloTag enabled target-specific PLP. For the Flag-tag strategy, we first transfected A549 cells with the Flag-EGFR construct and serum-starved the cells for >12 hours to remove serum-induced EGF signaling. An EY-conjugated Flag antibody, prepared as previously reported,^21^ was added to bind the extracellularly displayed Flag-tag, followed by addition of cell-penetrant photo-probes. Cells were then illuminated with blue light for 2 minutes. Flow cytometry confirmed that significant labeling was achieved using aryl-diazirine-biotin, aryl-azide-biotin and phenol-biotin with 2 min blue light activation (**Figure 1D**). Cells were lysed before biotinylated proteins were purified on streptavidin beads, trypsinized and analyzed by mass spectrometry (MS). EGFR was one of the most enriched proteins with all three photo-probes (**Figure S1C, S1D**).

To attach EY to the intracellular domain of the engineered EGFR-HaloTag, we synthesized a cell-penetrant EY hexyl-chloride derivative we call EY-HaloTag Ligand (EY-HTL) as previously reported (**Figure S2A**).^28^ Recent work has shown that conjugation using the HaloTag and HaloTag Ligand pair is highly efficient, selective, and compatible for cellular experiments.^29-31^ Similar to the Flag-tag eMultiMap workflow, we incubated the A549 cells expressing EGFR-HaloTag with EY-HTL to allow selective and covalent anchoring of the EGFR-HaloTag. To remove free EY-HTL from cells, we repeatedly washed the cells four times with fresh media including 15 min soaks between washes (**Figure S2A**). The labeling and washout procedure was optimized using a fluorophore with HaloTag Ligand developed by the Lavis lab, JF646-HTL.^32^ Flow cytometry showed dramatically increased fluorescent signal for the A549 cells expressing EGFR-HaloTag compared to A549 parental cells validating the washout procedure (**Figure S2B**). To test the specificity of the EY-HTL on a non-interacting target using this procedure, we transfected GFP into A549 cells with or without HaloTag and incubated with EY-HTL (**Figure S2C**). We conducted the iMultiMap experiments and observed selective enrichment of GFP over any other native protein for all three photo-probes (**Figure S2D**). We then performed the same workflow using EY-HTL on cells expressing EGFR-HaloTag and saw significant biotinylation (**Figure 2E**). These results confirmed that target-specific cellular labeling was achievable with genetically encoded Flag-tag and HaloTag and that they were compatible with the photocatalytic proximity labeling workflow.

**Figure 2:**
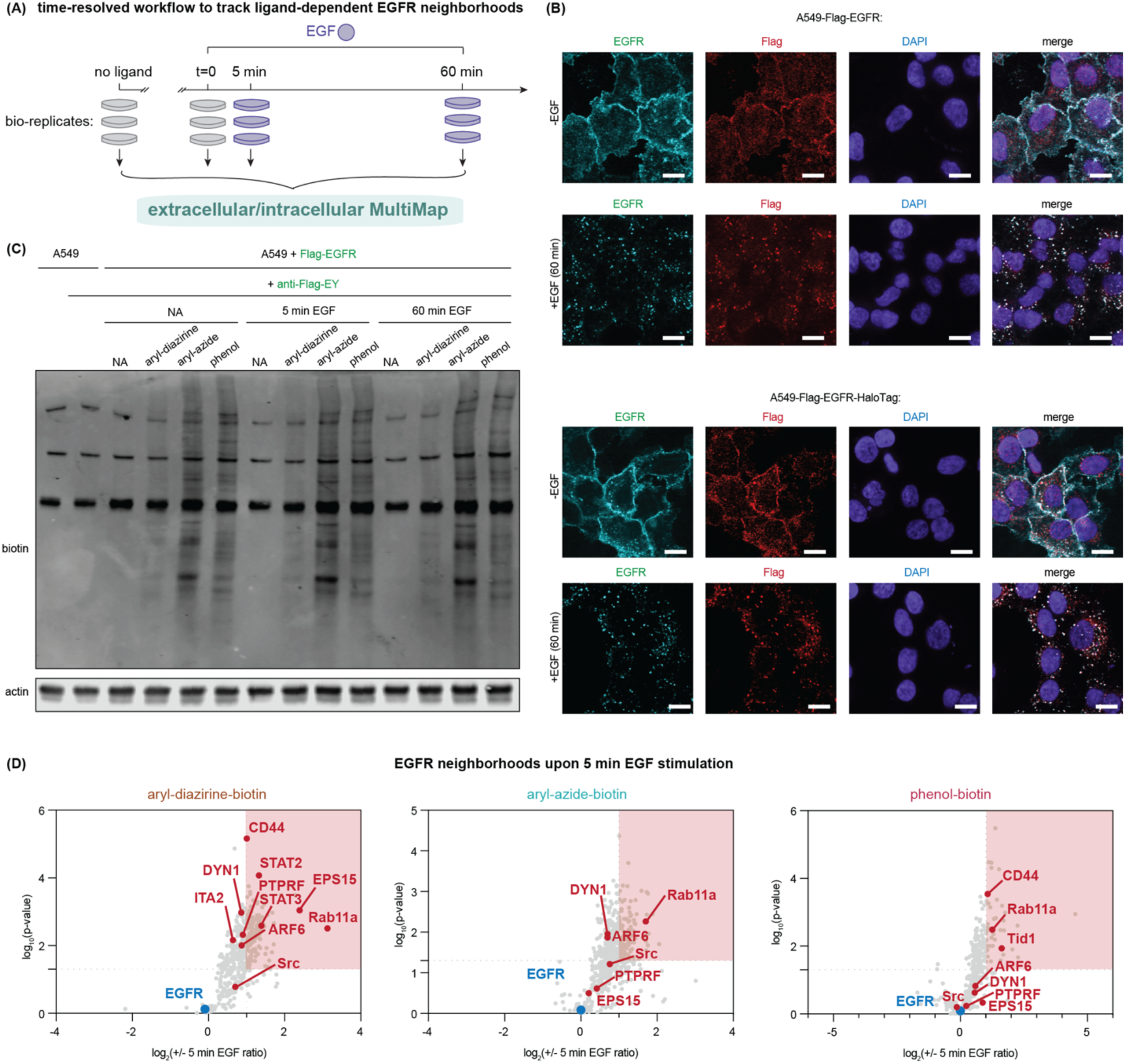
Rapid changes in EGFR neighborhoods upon EGF stimulation captured by time-resolved MultiMap. **(A)** General time-resolved MultiMap workflows to monitor EGF-activated EGFR neighborhood changes. **(B)** Confocal microscopy images of endogenous and Flag-EGFR (top) or EGFR-HaloTag (bottom) in cells treated with or without EGF. The staining of Flag-EGFR and EGFR-Halo overlapped those from the endogenous EGFR, suggesting that trafficking is indistinguishable. Scale bar = 20 μm. **(C)** WB analysis showing cellular biotinylation of A549 cells expressing Flag-EGFR using different photo-probes with short (5 min) and prolonged (60 min) EGF stimulation. **(D)** Volcano plots of rapid changes of on-state EGFR neighborhoods using eMultiMap. Labeled proteins using anti-Flag-EY were compared with or without 5 min EGF stimulation using aryl-diazirine-biotin, aryl-azide-biotin, phenol-biotin, respectively. Significantly enriched proteins are highlighted in red [log2(fold enrichment) ≥ 1, P < 0.05, unique peptide ≥ 2, n = 3] and listed in **Table S7-9**.

### Changes in the EGFR neighborhood upon 5 min treatment with EGF

Signaling through EGFR is known to be transient with an activation stage followed by a deactivation and turnover phase.^6, 7, 33^ We sought to monitor the changes of EGFR neighborhoods upon ligand activation in a time-resolved manner. EGF-triggered EGFR processes such as phosphorylation, dephosphorylation and internalization occur rapidly within minutes to over an hour. We found that even short light exposure (2 min) triggered ample protein labeling, making it possible to capture neighborhood changes with high temporal precision (**Figure 2A**).

We further showed via confocal imaging that Flag-EGFR and EGFR-HaloTag traffic with similar kinetics and localization as endogenous EGFR before and after EGF activation (**Figure 2B, S3A**). All proteins were largely internalized with 60 min EGF stimulation with an overall Pearson correlation coefficient above 0.6. We then performed the time-resolved Flag-tag eMultiMap workflow on cells in the presence of EGF (**Figure 2A**). Through WB analysis, we observed significant labeling on cells after EGF treatment for 5 min and 60 min, respectively (**Figure 2C**).

We then conducted large-scale proteomics experiments at different time-points by capturing biotinylated proteins on streptavidin beads, followed by trypsinization and MS analysis. We found a large number of proteins enriched upon 5 min EGF activation over the ligand-free state. After normalizing protein enrichment ratio by EGFR, we identified 158 proteins increased in labeling with aryl-diazirine-biotin, 149 proteins with aryl-azide-biotin and 87 proteins with phenol-biotin using the same statistical threshold [log2(fold enrichment) ≥ 1, P < 0.05, unique peptide ≥ 2, n = 3]. The volcano plot in **Figure 2D**, heatmap in **Figure S3B** and Venn diagram in **Figure S3C** showed overlap among the three probes. As expected from our previous observation,^21^ differential residue accessibility and labeling radius of each probe may contribute to the differences, suggesting that all probes are essential for a more holistic view of the neighborhood.

Among the neighboring proteins we identified, a number of candidates were reported to directly interact with EGFR or functionally associate with EGFR. For example, the Ras-related protein, Rab11a, is reported to co-localize with EGFR upon EGF stimulation^34^ and Rab11a overexpression accelerates EGFR recycling in cells.^35^ The EGFR pathway substrate EPS15, also named PHD3, was consistently found in our data and is reported to be associated with EGFR internalization and degradation.^36, 37^ We also found ARF6, an ADP-ribosylation factor, to be highly enriched upon treatment with EGF. ARF6 is reported to interact with EGFR and dependent on EGFR palmitoylation and ARF6 myristylation.^38^ Dynamin1 (DYN1) was highly enriched and is known to regulate EGFR internalization as demonstrated in a knock-out (KO) experiment.^39^ Other enriched targets with one or more photo-probes include the known functional interactors such as a tyrosine kinase protein Src,^40^ DnaJ homolog subfamily A member 3 (Tid1),^41^ the known transcription factor interactor STAT3,^42, 43^ the known extracellular stabilizer CD44,^44-46^ as well as potential regulator tyrosine-protein phosphatase transmembrane receptor PTPRF.^47^

### Changes in EGF-stimulated EGFR neighbors over 60 min treatment

Next, we sought to monitor the dynamics of these protein-protein interactions over prolonged EGF stimulation and evaluate whether the EGFR neighbors still remain proximal to EGFR. We performed the same proteomics workflow and profiled the protein neighbors of EGFR upon 60 min EGF incubation (**Figure 3A**). We found that significantly fewer proteins were enriched at the 60 min time-point with all three probes: 76 proteins enriched with aryl-diazirine-biotin, 36 proteins enriched with aryl-azide-biotin, 24 enriched with phenol-biotin using the same statistical threshold [log2(fold enrichment) ≥ 1, P < 0.05, unique peptide ≥ 2, n = 3] (volcano plot **Figure 3A**, heatmap **Figure S4A**). Protein hits enriched at 60 min also showed overlap among three different probes (**Figure S4B**). Some that overlap with 5 min incubation (**Figure S4C**) include Rab11a, EPS15, ARF6, DYN1 and Src (**Figure 3A**).

**Figure 3:**
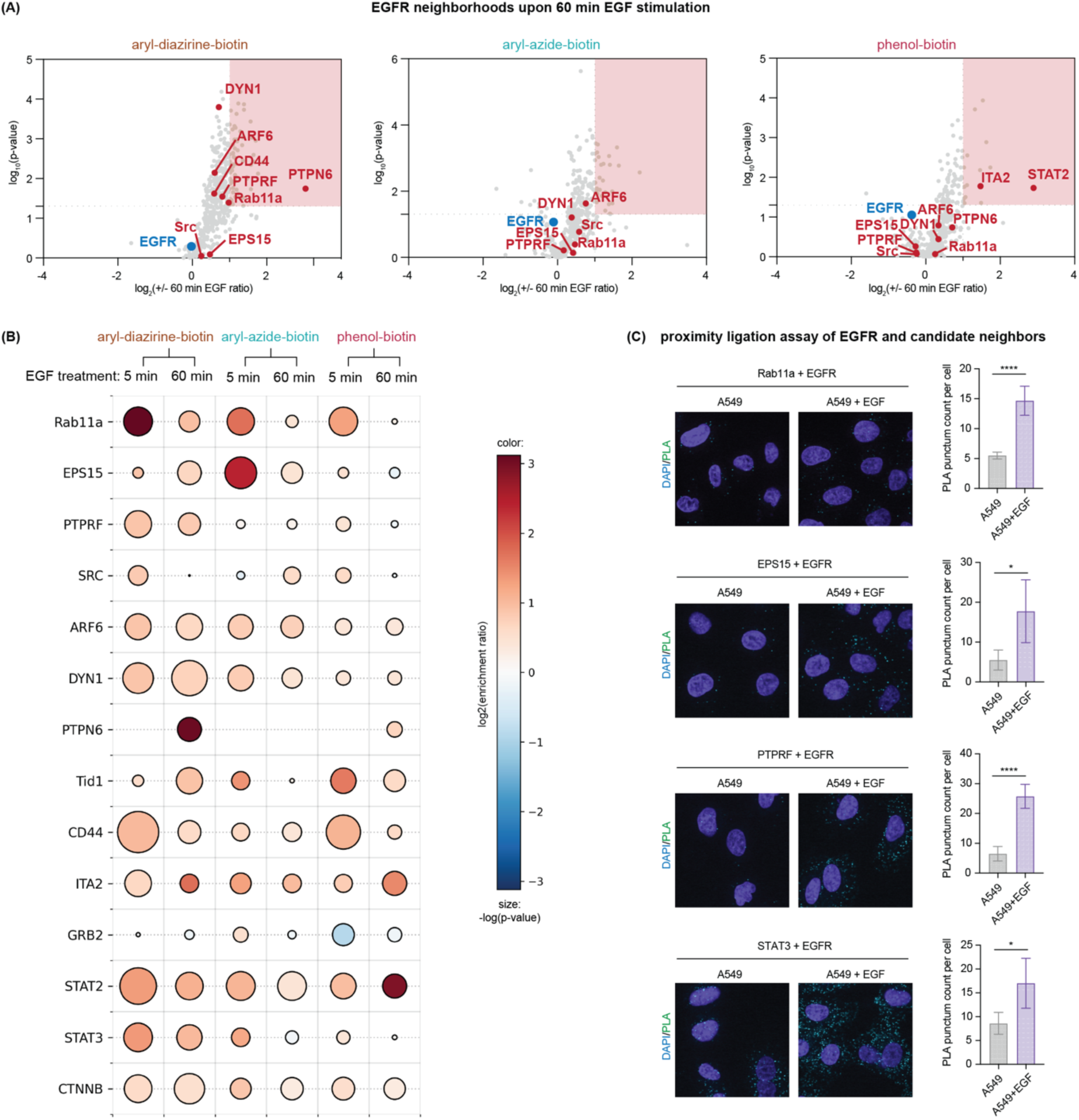
Time-resolved labeling by eMultiMap reflects kinetics of EGF-dependent EGFR neighbors. **(A)** Volcano plots of prolonged changes of on-state EGFR neighborhoods using eMultiMap. Biotinylated proteins using anti-Flag-EY were compared with or without 60 min of EGF stimulation using aryl-diazirine-biotin, aryl-azide-biotin, or phenol-biotin photo-probes. Significantly enriched proteins are highlighted in red [log2(fold enrichment) ≥ 1, P < 0.05, unique peptide ≥ 2, n = 3] and listed in **Table S10-12**. **(B)** Enrichment profiles of candidate EGFR neighbors at 5 min and 60 min EGF stimulation using the three photo-probes with different labeling range. Color (blue to red) represents increasing log2(fold enrichment) at indicated treatment time versus no EGF treatment. Size of bubbles indicates -log(p-value) across replicates with larger sizes indicating higher confidence. **(C)** Proximity ligation assay (PLA) confirmed colocalization of candidate EGFR neighbors with EGFR in an EGF-dependent manner. Scale bar = 20 μm. Data are represented as mean ± SD.

We compared the time-dependent enrichment for the top protein hits that were highly enriched in one of the datasets or enriched throughout all datasets via a balloon plot (**Figure 3B**). Rab11a, one of the most enriched proteins with 5 min EGF stimulation, was found to be highly enriched throughout all three probes at 5 min activation, However, Rab11A was enriched at significantly lower enrichment ratios with 60 min EGF stimulation. This suggests rapid association followed by disassociation and is consistent with the transient role Rab11A may have in trafficking. Similarly, EPS15 was highly enriched at 5 min using aryl-azide-biotin, but the enrichment ratio significantly dropped at 60 min.

Other proteins showed sustained levels of enrichment upon EGF stimulation such as PTPRF, Src, ARF6, DYN1, CD44 at 60 min when compared with enrichment level at 5 min (**Figure 3B**). Transcription factors such as STAT3, STAT2 and β-catenin (CTNNB) also showed little change in enrichment at 5 min and 60 min. Interestingly, enrichment ratios of several proteins became more prominent at 60 min compared with 5 min. This group of proteins included protein tyrosine phosphatase non-receptor type 6, PTPN6, also called SHP1, which has been reported to physically interact with EGFR,^48, 49^ as well as integrin A2 (ITA2). It is likely that these proteins are recruited later to EGFR.

To validate these candidate neighbors and confirm that their interaction with EGFR is dependent on ligand activation, we sought to visualize protein localization in situ using proximity ligation assays (PLA) (**Figure 3C).** PLA provides spatial proximity information with high sensitivity and is orthogonal to other assays such as co-immunoprecipitation.^50, 51^ We performed the PLA on five top hits, Rab11a, EPS15, PTPRF, STAT3 and Src (**Figure 3C**, **S4D**). When comparing ligand-free A549 cells and cells with EGF treatment, we found significant increases of PLA foci per cell upon EGF activation with Rab11a, EPS15, PTPRF and STAT3. These results orthogonally support the co-localization of these targets to EGFR upon EGF stimulation.

### iMultiMap enables both identification and validation of intracellular partners of EGFR

Most signaling receptors have an intracellular domain to transfer the binding information from the cell surface to intracellular signaling. Although the photo-probes we use here are cell permeable, we hypothesized that iMultiMap would be closer to events on the inside of cells and additionally serve as an orthogonal validation method for targets identified by eMultiMap. We performed a similar workflow labeling proteins in proximity of EGFR-HaloTag upon 5 min and 60 min EGF activation (**Figure 4A**). Cells expressing EGFR-HaloTag were serum starved and incubated with EY-HTL followed by a repeated washout step to remove non-covalently bound EY-HTL (**Figure S2, S5A**). EGF was added to cells before the aryl-diazirine-, aryl-azide- and phenol-biotin photo-probes were added and cells were illuminated with blue light for 2 min to induce biotinylation (**Figure S5A)**. Not surprisingly, we observed comparable bulk labeling in cells with no ligand, with 5 min EGF activation and 60 min EGF activation by WB analysis that reflects global biotinylation (**Figure 4A**).

**Figure 4:**
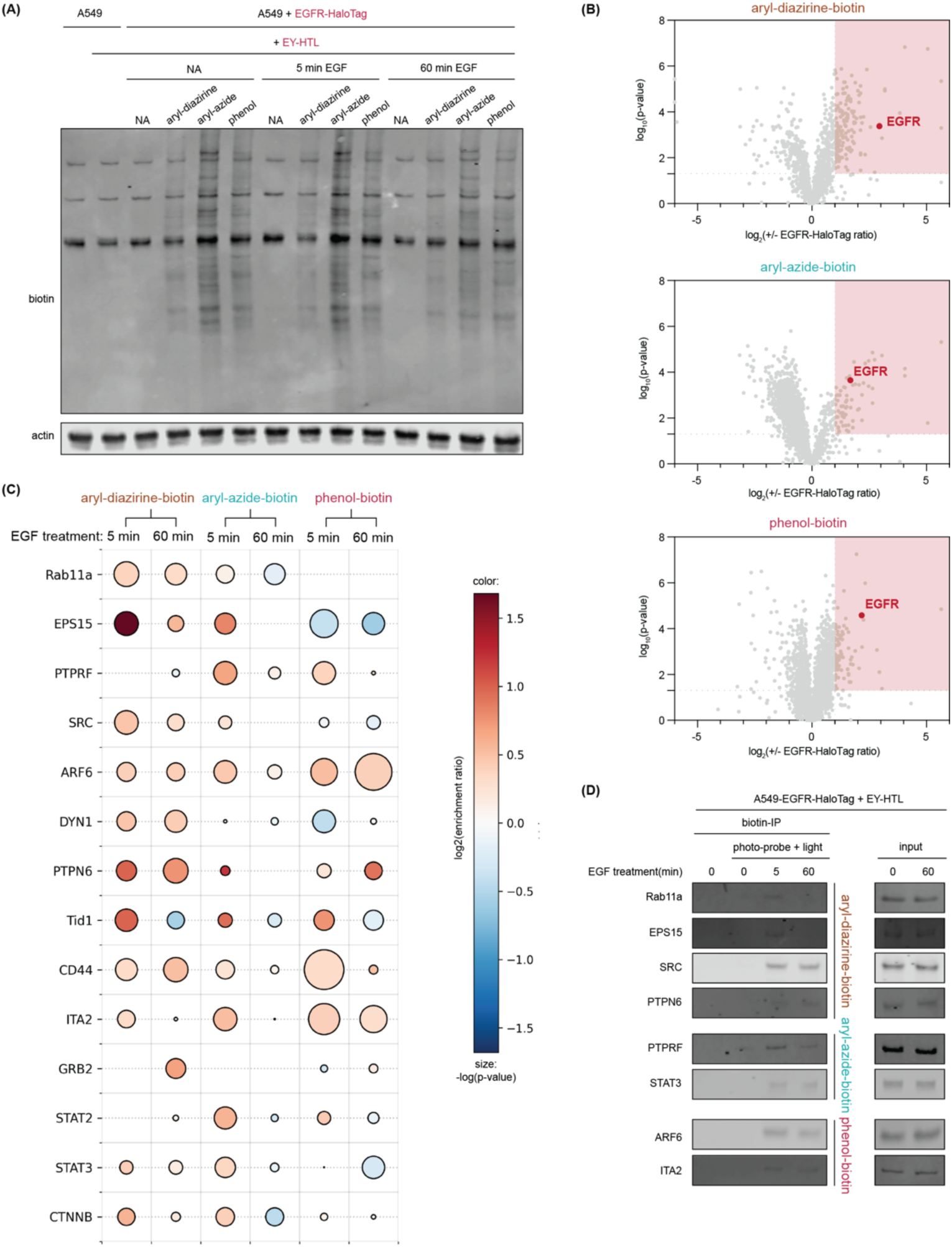
iMultiMap enabled identification and validation of intracellular neighbors of EGFR in an EGF-dependent manner. **(A)** Cellular labeling of A549 cells expressing EGFR-HaloTag using different photo-probes with short (5 min) and prolonged (60 min) EGF stimulation. **(B)** Volcano plots showing rapid changes of on-state EGFR neighborhoods using iMultiMap. Labeled proteins using EY-HTL were compared with and without 5 min EGF stimulation using the aryl-diazirine-biotin, aryl-azide-biotin, phenol-biotin photo-probes. Significantly enriched proteins are highlighted in red [log2(fold enrichment) ≥ 1, P < 0.05, unique peptide ≥ 2, n = 3] and listed in **Table S13-15**. **(C)** Enrichment profiles of candidate EGFR neighbors at 5 min and 60 min EGF stimulation using three photo-probes. Color (blue to red) represents increasing log2(fold enrichment) at indicated treatment time versus no EGF treatment. Size of bubbles indicates - log(p-value) across replicates with larger sizes indicating higher confidence. **(D)** Biotin-IP western blots confirm that EGFR neighbors are captured in situ with EY-HTL-induced iMultiMap at different time-points.

We then used our established proteomics workflow to identify the specific protein enriched (**Figure 4B**). We observed that EGFR was as one of the most enriched proteins in the datasets for all three probes comparing A549-WT and A549-EGFR-HaloTag cells (**Figure 4B**). We then treated cells with EGF and compared enriched proteins at the 5 min (**Figure S5B**) and 60 min (**Figure S5C**). At the 5 min time-point, we found 105 proteins enriched with aryl-diazirine-biotin, 59 proteins with aryl-azide-biotin and 229 proteins with phenol-biotin using the threshold [log2(fold enrichment) ≥ 1, P < 0.05, unique peptide ≥ 2, n = 3], respectively. At the 60 min time-point, we found 112 proteins enriched with aryl-diazirine-biotin, 13 proteins with aryl-azide-biotin and 25 proteins with phenol-biotin using the threshold [log2(fold enrichment) ≥ 1, P < 0.05, unique peptide ≥ 2, n = 3], respectively.

The individual protein time-dependent enrichment profiles from iMultiMap shared similarities with the ones from eMultiMap as visualized with balloon plots in **Figure 4C and 3B**, respectively. In particular, proteins such as Rab11a and EPS15 were significantly more enriched at the 5 min time-point than at the 60 min time-point. Most other proteins shared similar enrichment patterns from those from eMultiMap. However, we did find some differences in these time-dependent profiles between eMultiMap and iMultiMap. For example, Tid1 demonstrated more significant differences of 5 min and 60 min enrichment than the ones quantified through eMultiMap. This is not surprising since Tid1 is an intracellular mitochondrial-associated protein that reportedly regulates EGF-stimulated interaction with HSP70 chaperone system.^41^ PTPN6, or namely SHP-2, on the other hand, was similarly more enriched at 60 min by iMultiMap using phenol-biotin, possibly for the same reason. Nonetheless, in general the enrichment was comparable between the two datasets of eMultiMap and iMultiMap, suggesting that the two methods can provide reciprocal validation to each other.

We further used WB analysis to visualize the specific biotinylated proteins enriched through iMultiMap (**Figure 4D**). We confirmed the increase in biotinylation of Rab11a, EPS15, Src and PTPN6 with aryl-diazirine-biotin enrichment, as well as PTPRF, STAT3 with aryl-azide-biotin enrichment, and ARF6, ITA2 with phenol-biotin enrichment. Remarkably, the enrichment blots were also capable of differentiating labeling levels at different time-points of EGF ligand activation. Enriched Rab11a and EPS15, for example, were higher in amount at 5 min EGF activation time-point than at 60 min EGF activation time-point. These data provide an orthogonal validation to the PLP, and support that iMultiMap is complementary to eMultiMap for neighborhood identification.

### EGFR neighbors are associated with EGF-activated EGFR functions

EGFR is known to undergo staged signaling events over an hour or so upon EGF ligand activation, including phosphorylation, internalization, degradation and transcriptional regulation.^7, 52^ We were curious if any of the targets we identified by eMultiMap and iMultiMap were regulating these processes. To study this, we genetically manipulated selected proteins of interest and monitored the corresponding EGFR responses upon EGF activation (**Figure 5**).

**Figure 5:**
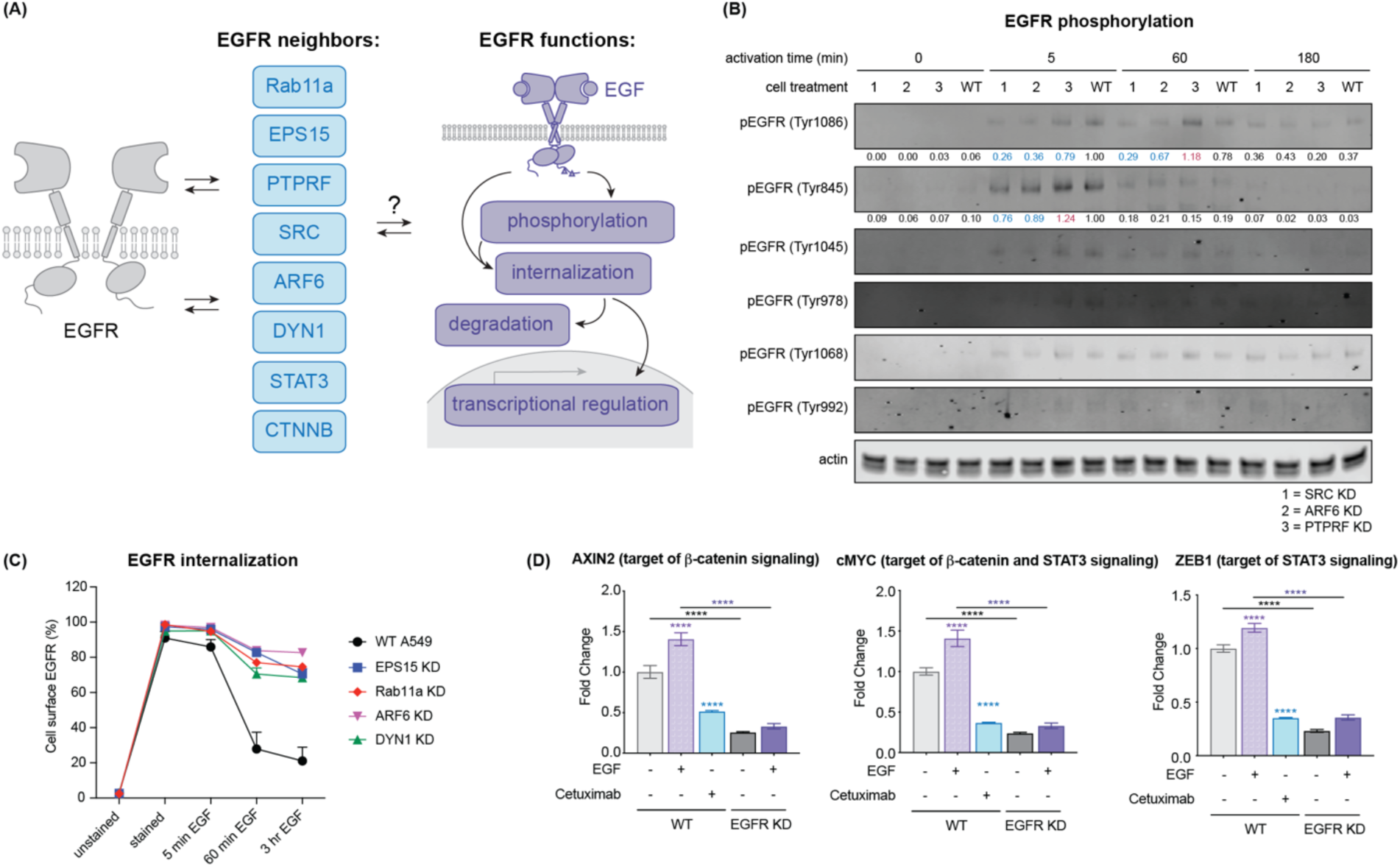
Dynamic association of EGF-activated EGFR neighbors lead to functional regulation of EGFR. **(A)** Summary of dynamically associating EGFR neighbors that change most with time as viewed by eMultiMap and iMultiMap. **(B)** Phosphorylation of EGFR at known sites in A549 cells with and without neighbor gene knock-downs. **(C)** EGFR internalization in A549 cells monitored with and without neighbor gene knock-downs. **(D)** EGFR regulation on transcriptional activities of STAT3 and β-catenin measured by qRT-PCR analysis. Data are represented as mean ± SD.

Phosphorylation is an immediate and critical regulatory mechanism behind EGFR signaling cascades. Thus, we first examined various phosphorylation sites on EGFR, including Tyr1086, Tyr845, Tyr1045, Tyr978, Tyr1068 and Tyr992 sites, which are known to be associated with different functions of EGFR such as signaling and degradation (**Figure 5B**, **S6A**).^33, 53^ In WT A549 cells, we confirmed known differential kinetic patterns of phosphorylation at different sites. For example, we observed known phosphorylation events on the Tyr1045, Tyr 845, Tyr978, Tyr992 sites that returned to basal levels after 1 hour of EGF stimulation, while prolonged phosphorylation at Tyr1086, Tyr 1068 sites showed sustained response to EGF stimulation. With Src or ARF6 knocked down, lower levels of EGFR phosphorylation were observed for Tyr1086 and Tyr845 sites with mild influence on the Tyr1068 site phosphorylation, a major autophosphorylation site involved in the MAPK signaling cascade. Interestingly, knock-down of the PTPRF phosphatase, a known tumor suppressor, led to increased phosphorylation of the early Tyr845 and late Tyr1086 sites while the other phospho-sites were largely unaffected.

We next sought to track time-dependent EGFR internalization by monitoring cell-surface EGFR at different time-points of EGF activation (**Figure 5C**, **S6B-C**). With wild-type A549 cells, EGFR was internalized rapidly starting from 5 min EGF activation and was kept internalized at the 60 min and 3 hour time-points (**Figure S6B**). With decreased presence of Rab11a, EPS15, DYN1 and ARF6, cell surface EGFR internalization was greatly inhibited (**Figure 5C**, **S6C-D**). After three hours of EGF stimulation >60% EGFR was internalized in wild-type A549 cells. When Rab11a, EPS15, DYN1 or ARF6 were individually knocked down, we observed only ∼20% EGFR internalization at the 3 hour time-point. These data suggest that Rab11a, EPS15, DYN1 and ARF6 significantly contribute to EGFR internalization upon EGF activation.

After internalization, EGFR can be degraded which regulates EGFR availability and activity. By knocking down EPS15 and DYN1 (**Figure S6C, S6E**), we observed significant reduction in EGFR degradation compared to that in WT cells, which extended to a 16 hour time-point (**Figure S6E**). When knocking down Rab11a and ARF6, while no significant changes were observed at the 1 hour and 3 hour time-points, EGFR degradation was enhanced at the 16 hour time-point. This result was possibly due to the additional roles of these neighbors such as Rab11a being expected to mediate EGFR recycling from endosome back to membrane.^34, 54^ ARF6 additionally regulates EGFR level by engaging the EGFR sorting process from Golgi to membrane, which is possibly the cause of this differential EGFR level at the prolonged time-point.^38^ These data support the roles of these proteins in promoting trafficking ultimately leading to EGFR turnover and homeostasis.

Activated EGFR also drives transcription through the regulation of a number of transcription factors including STAT3 and β-catenin.^42, 43, 55, 56^ As expected, both STAT3 and β-catenin were enriched through our eMultiMap and iMultiMap PLP labeling upon EGF stimulation, indicating that they can physically interact (**Figure 3B**, **4C**). To further confirm the regulatory function of EGFR in the A549 cells, we conducted EGFR knock-down and monitored STAT3 and β-catenin activities via qPCR analysis (**Figure 5D**, **S6F**). We observed increased activity of STAT3 and β-catenin upon EGF stimulation, leading to increased level of the β-catenin-specific target gene AXIN2, the STAT3-specific target gene ZEB1, and the shared β-catenin/STAT3 target gene cMYC.^57, 58^ EGFR inhibition with cetuximab on the other hand, reduced the gene expression for both systems. When EGFR is knocked down, RNA levels of all these genes were significantly reduced and minimal induction was observed in the presence of EGF, sharing the pattern with EGFR inhibition. These results support that transcriptional activities of both STAT3 and β-catenin are modulated by EGFR activity, establishing them as functional neighbors of EGFR (**Figure 6**).

**Figure 6:**
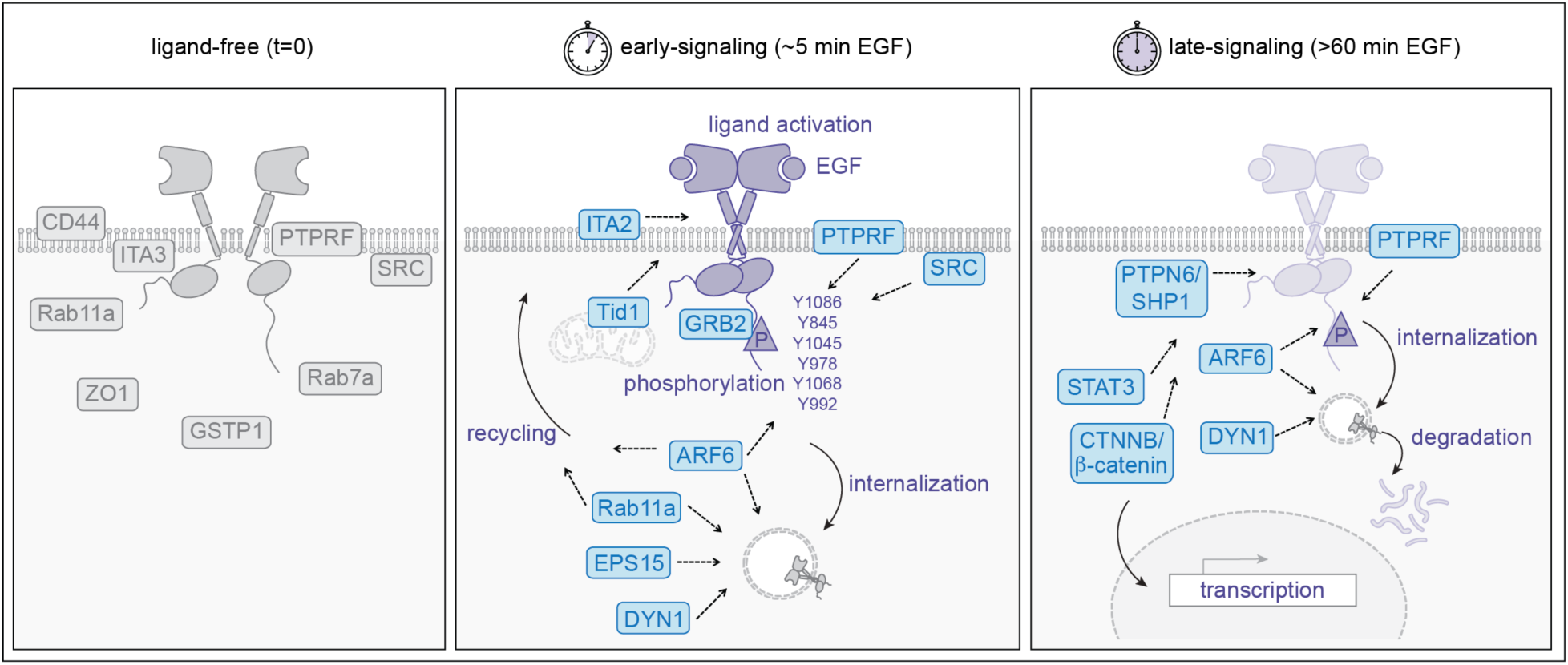
Model of EGFR association with functional neighbors identified via spatiotemporally resolved MultiMap. We mapped the dynamic neighborhoods of EGFR upon ligand activation via the dual set of eMultiMap and iMultiMap, which uncovered the time-dependent association of these functional neighbors with each critical EGFR signaling processes including phosphorylation, internalization, degradation and transcriptional activation.

## Discussion

We present a set of new spatiotemporally resolved MultiMap workflows, eMultiMap and iMultiMap, that enable temporal tracking of protein neighborhoods during EGF-induced signaling processes. Others as well as ourselves have previously demonstrated labeling strategies to attach the photocatalyst using a selective antibody to the protein of interest (POI), such as cetuximab,^21^ receptor-specific ligands or protein-binding small molecules.^59-64^ However, these approaches utilize epitopes on the POI which can affect protein neighbors. By appending genetically encoded tags on either side of the POI, in our case EGFR, we successfully introduced our photocatalyst, Eosin Y, without intentionally blocking potential neighborhood interactions. Moreover, this genetically encoded ectopic tagging strategy does not require specific antibodies to the POI, thus greatly expanding the targetable proteome for photocatalytic PLP. Given there may be cases where the ectopic tag could impact functions of the POI, it is important to evaluate if the ectopic tag has a significant functional impact in each use case.

By visualizing EGFR neighborhoods from both outside and inside the cells via eMultiMap and iMultiMap, we obtained orthogonal and complementary datasets via proteomics. The majority of hits were shared between the two datasets, which is not surprising given that photo-probes can diffuse bi-directionally through the cell membrane. We did however observe targets specific to eMultiMap and iMultiMap, as well as targets with differential enrichment levels. This most likely derives from the localization of these targets or general accessibility for photo-probes in these specific cellular milleu. For example, once EGFR is internalized, one can imagine that iMultiMap will have an advantage over eMultiMap. Additionally, we observed significantly constrained level of labeling for phenol-biotin for iMultiMap compared to eMultiMap. The phenoxy radicals have the longest half-life and thus expected to have the longest distance of labeling. The more confined range of the phenol-biotin we observed in iMultiMap is possibly due to the presence of radical quenching molecules or enzymes within live cells not found on the outside. Further characterization of the intracellular labeling environment will be useful to better understand the labeling radii and quenching mechanisms that may reside inside cells.

It is known that EGFR activation undergoes a series of time-staged signaling processes. We believe eMultiMap and iMultiMap are well suited to probe the dynamic neighborhoods given the robustness of labeling within 2-5 min illumination window, allowing us to visualize them with enhanced temporal resolution. This may be applicable to other proteins or proteoforms during their active signaling, including but not limited to aberrant mutants,^65, 66^ drug mechanisms,^67, 68^ and cell-/organ-specific neighborhoods.^69, 70^

By performing this spatiotemporally resolved MultiMap workflow, we identified functional targets and confirmed their association to EGFR functions that span through a wide range of timescales. With direct EGFR phosphorylation happening within minutes upon EGF stimulation, EGFR internalization and degradation happens in hours followed by nuclear translocation that leads to EGFR regulation of transcriptional activities. We were able to systematically track functional neighbors in each process, including SRC, ARF6 and PTPRF during their regulation of EGFR phosphorylation, Rab11a, EPS15, DYN1 and ARF6 when regulating EGFR internalization as well as STAT3 and β-catenin as downstream transcription factors, which provided a list of targetable entities (**Figure 6**). For example, PTPRF, PTPN6 and EPS15 are known serve as tumor suppressors,^71-74^ whereas Src kinase, Rab11a and CD44 may be tumor promoters.^46, 75-77^ We envision this mechanistically detailed proximity map of EGFR functional neighborhood will help dissect the mechanisms behind these different processes and facilitate phenotype-specific probing of EGFR.

Taken together, the spatiotemporally resolved MultiMap workflows capture interactome snapshots with high resolution, allowing insights to be gleaned holistically both on the cell surface and within the cells. By dissecting the dynamic interactions of cell surface proteins during its active signaling cascade, we can visualize the horizontal signaling of EGFR that occurs simultaneously upon vertical ligand activation. We believe such studies will reveal new functional interaction and lead to new targets for therapeutic development.

## Acknowledgements

We thank Kevin Leung (UCSF), Bo Huang (UCSF), Klaus Yserentant (UCSF), Danielle Swaney (UCSF) and Kari Herrington (Center for Advanced Light Microscopy, UCSF) for insightful discussions and technical support. We also thank Luke Lavis (Janelia) for providing HaloTag-JaneliaFluor compounds as well as Alice Cheng (UCSF) for her experimental assistance. We are grateful to generous support from NIH-1R01CA248323-01 (J.A.W.; Z.L.), NIH-R35GM122451 (J.A.W., Z.L.), the Anne Wojcicki Foundation (J.A.W.), the Hind Professorship in Pharmaceutical Sciences (J.A.W.), UCSF Mary Anne Koda-Kimble Seed Award for Innovation (Z.L.), Natural Sciences and Engineering Research Council of Canada Postdoctoral Fellowship PDF-578176-2023 (W.N.), James B. Pendleton Charitable Trust (W.N.; J.A.D.), Gladstone Institutes (W.N.; J.A.D.), National Heart, Lung, and Blood Institute grant 1R21HL173710-01 (J.A.D.), Lawrence Livermore National Labs PROTECT DE-AC52-07NA27344 (J.A.D.), NIH-U54CA224081 (T.G.B.), NIH-U01CA217882 (T.G.B.), R01CA204302 (T.G.B.), R01CA211052 (T.G.B.), Chan-Zuckerberg Biohub (T.G.B.), NIGMS R35GM134948 (Y.J.), NIH-DP5OD036136 (K.J.S.) and the UCSF Sandler Fellowship (K.J.S.). J.A.D. is a Howard Hughes Medical Institute Investigator.

## Author contributions

Z.L. and J.A.W. designed the project, analyzed the data, and wrote the manuscript, with input from all authors. W.N. and J.A.D. generated and optimized gene knockdown. H.W., Y.J. and Z.L. performed confocal images and analyzed data. Y-T.C. and T.G.B. designed and performed RNA quantification. Z.L. and K.J.S. generated plasmids and purified proteins.

## Supplementary information

Figures S1-S6.

Tables S1-S21.

Methods.

## References

1. Lemmon, M. A.; Schlessinger, J., Cell signaling by receptor tyrosine kinases. Cell 2010, 141 (7), 1117–34.

2. Ogiso, H.; Ishitani, R.; Nureki, O.; Fukai, S.; Yamanaka, M.; Kim, J. H.; Saito, K.; Sakamoto, A.; Inoue, M.; Shirouzu, M.; Yokoyama, S., Crystal structure of the complex of human epidermal growth factor and receptor extracellular domains. Cell 2002, 110 (6), 775–87.

3. Ferguson, K. M.; Berger, M. B.; Mendrola, J. M.; Cho, H. S.; Leahy, D. J.; Lemmon, M. A., EGF activates its receptor by removing interactions that autoinhibit ectodomain dimerization. Mol Cell 2003, 11 (2), 507–17.

4. Jura, N.; Endres, N. F.; Engel, K.; Deindl, S.; Das, R.; Lamers, M. H.; Wemmer, D. E.; Zhang, X.; Kuriyan, J., Mechanism for activation of the EGF receptor catalytic domain by the juxtamembrane segment. Cell 2009, 137 (7), 1293–307.

5. Kaplan, M.; Narasimhan, S.; de Heus, C.; Mance, D.; van Doorn, S.; Houben, K.; Popov-Celeketic, D.; Damman, R.; Katrukha, E. A.; Jain, P.; Geerts, W. J. C.; Heck, A. J. R.; Folkers, G. E.; Kapitein, L. C.; Lemeer, S.; van Bergen En Henegouwen, P. M. P.; Baldus, M., EGFR Dynamics Change during Activation in Native Membranes as Revealed by NMR. Cell 2016, 167 (5), 1241–1251 e11.

6. Reddy, R. J.; Gajadhar, A. S.; Swenson, E. J.; Rothenberg, D. A.; Curran, T. G.; White, F. M., Early signaling dynamics of the epidermal growth factor receptor. Proc Natl Acad Sci U S A 2016, 113 (11), 3114–9.

7. Freed, D. M.; Bessman, N. J.; Kiyatkin, A.; Salazar-Cavazos, E.; Byrne, P. O.; Moore, J. O.; Valley, C. C.; Ferguson, K. M.; Leahy, D. J.; Lidke, D. S.; Lemmon, M. A., EGFR Ligands Differentially Stabilize Receptor Dimers to Specify Signaling Kinetics. Cell 2017, 171 (3), 683–695 e18.

8. Maemondo, M.; Inoue, A.; Kobayashi, K.; Sugawara, S.; Oizumi, S.; Isobe, H.; Gemma, A.; Harada, M.; Yoshizawa, H.; Kinoshita, I.; Fujita, Y.; Okinaga, S.; Hirano, H.; Yoshimori, K.; Harada, T.; Ogura, T.; Ando, M.; Miyazawa, H.; Tanaka, T.; Saijo, Y.; Hagiwara, K.; Morita, S.; Nukiwa, T.; North-East Japan Study, G., Gefitinib or chemotherapy for non-small-cell lung cancer with mutated EGFR. N Engl J Med 2010, 362 (25), 2380–8.

9. Nakai, K.; Hung, M. C.; Yamaguchi, H., A perspective on anti-EGFR therapies targeting triple-negative breast cancer. Am J Cancer Res 2016, 6 (8), 1609–23.

10. Mok, T. S.; Wu, Y. L.; Thongprasert, S.; Yang, C. H.; Chu, D. T.; Saijo, N.; Sunpaweravong, P.; Han, B.; Margono, B.; Ichinose, Y.; Nishiwaki, Y.; Ohe, Y.; Yang, J. J.; Chewaskulyong, B.; Jiang, H.; Duffield, E. L.; Watkins, C. L.; Armour, A. A.; Fukuoka, M., Gefitinib or carboplatin-paclitaxel in pulmonary adenocarcinoma. N Engl J Med 2009, 361 (10), 947–57.

11. Soria, J. C.; Ohe, Y.; Vansteenkiste, J.; Reungwetwattana, T.; Chewaskulyong, B.; Lee, K. H.; Dechaphunkul, A.; Imamura, F.; Nogami, N.; Kurata, T.; Okamoto, I.; Zhou, C.; Cho, B. C.; Cheng, Y.; Cho, E. K.; Voon, P. J.; Planchard, D.; Su, W. C.; Gray, J. E.; Lee, S. M.; Hodge, R.; Marotti, M.; Rukazenkov, Y.; Ramalingam, S. S.; Investigators, F., Osimertinib in Untreated EGFR-Mutated Advanced Non-Small-Cell Lung Cancer. N Engl J Med 2018, 378 (2), 113–125.

12. Guardiola, S.; Varese, M.; Sanchez-Navarro, M.; Giralt, E., A Third Shot at EGFR: New Opportunities in Cancer Therapy. Trends Pharmacol Sci 2019, 40 (12), 941–955.

13. Brand, T. M.; Iida, M.; Wheeler, D. L., Molecular mechanisms of resistance to the EGFR monoclonal antibody cetuximab. Cancer Biol Ther 2011, 11 (9), 777–92.

14. Sequist, L. V.; Waltman, B. A.; Dias-Santagata, D.; Digumarthy, S.; Turke, A. B.; Fidias, P.; Bergethon, K.; Shaw, A. T.; Gettinger, S.; Cosper, A. K.; Akhavanfard, S.; Heist, R. S.; Temel, J.; Christensen, J. G.; Wain, J. C.; Lynch, T. J.; Vernovsky, K.; Mark, E. J.; Lanuti, M.; Iafrate, A. J.; Mino-Kenudson, M.; Engelman, J. A., Genotypic and histological evolution of lung cancers acquiring resistance to EGFR inhibitors. Sci Transl Med 2011, 3 (75), 75ra26.

15. Quinn, P.; Griffiths, G.; Warren, G., Density of newly synthesized plasma membrane proteins in intracellular membranes II. Biochemical studies. J. Cell Biol. 1984, 98 (6), 2142–7.

16. Liang, Y.; Fotiadis, D.; Filipek, S.; Saperstein, D. A.; Palczewski, K.; Engel, A., Organization of the G protein-coupled receptors rhodopsin and opsin in native membranes. J Biol Chem 2003, 278 (24), 21655–21662.

17. Jacobson, K.; Mouritsen, O. G.; Anderson, R. G., Lipid rafts: at a crossroad between cell biology and physics. Nat. Cell Biol. 2007, 9 (1), 7–14.

18. Huang, W. Y. C.; Boxer, S. G.; Ferrell, J. E., Membrane localization accelerates association under conditions relevant to cellular signaling. Proc. Natl. Acad. Sci. 2024, 121 (10), e2319491121.

19. Geri, J. B.; Oakley, J. V.; Reyes-Robles, T.; Wang, T.; McCarver, S. J.; White, C. H.; Rodriguez-Rivera, F. P.; Parker, D. L., Jr.; Hett, E. C.; Fadeyi, O. O.; Oslund, R. C.; MacMillan, D. W. C., Microenvironment mapping via Dexter energy transfer on immune cells. Science 2020, 367 (6482), 1091–1097.

20. Bartholow, T. G.; Burroughs, P. W. W.; Elledge, S. K.; Byrnes, J. R.; Kirkemo, L. L.; Garda, V.; Leung, K. K.; Wells, J. A., Photoproximity Labeling from Single Catalyst Sites Allows Calibration and Increased Resolution for Carbene Labeling of Protein Partners In Vitro and on Cells. ACS Cent Sci 2024, 10 (1), 199–208.

21. Lin, Z.; Schaefer, K.; Lui, I.; Yao, Z.; Fossati, A.; Swaney, D. L.; Palar, A.; Sali, A.; Wells, J. A., Multiscale photocatalytic proximity labeling reveals cell surface neighbors on and between cells. Science 2024, 385 (6706), eadl5763.

22. Oakley, J. V.; Buksh, B. F.; Fernandez, D. F.; Oblinsky, D. G.; Seath, C. P.; Geri, J. B.; Scholes, G. D.; MacMillan, D. W. C., Radius measurement via super-resolution microscopy enables the development of a variable radii proximity labeling platform. Proc Natl Acad Sci U S A 2022, 119 (32), e2203027119.

23. Li, S.; Schmitz, K. R.; Jeffrey, P. D.; Wiltzius, J. J.; Kussie, P.; Ferguson, K. M., Structural basis for inhibition of the epidermal growth factor receptor by cetuximab. Cancer Cell 2005, 7 (4), 301–11.

24. Yonesaka, K.; Zejnullahu, K.; Okamoto, I.; Satoh, T.; Cappuzzo, F.; Souglakos, J.; Ercan, D.; Rogers, A.; Roncalli, M.; Takeda, M.; Fujisaka, Y.; Philips, J.; Shimizu, T.; Maenishi, O.; Cho, Y.; Sun, J.; Destro, A.; Taira, K.; Takeda, K.; Okabe, T.; Swanson, J.; Itoh, H.; Takada, M.; Lifshits, E.; Okuno, K.; Engelman, J. A.; Shivdasani, R. A.; Nishio, K.; Fukuoka, M.; Varella-Garcia, M.; Nakagawa, K.; Janne, P. A., Activation of ERBB2 signaling causes resistance to the EGFR-directed therapeutic antibody cetuximab. Sci Transl Med 2011, 3 (99), 99ra86.

25. Uhlen, M.; Oksvold, P.; Fagerberg, L.; Lundberg, E.; Jonasson, K.; Forsberg, M.; Zwahlen, M.; Kampf, C.; Wester, K.; Hober, S.; Wernerus, H.; Bjorling, L.; Ponten, F., Towards a knowledge-based Human Protein Atlas. Nat Biotechnol 2010, 28 (12), 1248–50.

26. Wisniewski, J. R.; Hein, M. Y.; Cox, J.; Mann, M., A “proteomic ruler” for protein copy number and concentration estimation without spike-in standards. Mol Cell Proteomics 2014, 13 (12), 3497–506.

27. Jorissen, R. N.; Walker, F.; Pouliot, N.; Garrett, T. P.; Ward, C. W.; Burgess, A. W., Epidermal growth factor receptor: mechanisms of activation and signalling. Exp Cell Res 2003, 284 (1), 31–53.

28. Takemoto, K.; Matsuda, T.; McDougall, M.; Klaubert, D. H.; Hasegawa, A.; Los, G. V.; Wood, K. V.; Miyawaki, A.; Nagai, T., Chromophore-assisted light inactivation of HaloTag fusion proteins labeled with eosin in living cells. ACS Chem Biol 2011, 6 (5), 401–6.

29. Los, G. V.; Encell, L. P.; McDougall, M. G.; Hartzell, D. D.; Karassina, N.; Zimprich, C.; Wood, M. G.; Learish, R.; Ohana, R. F.; Urh, M.; Simpson, D.; Mendez, J.; Zimmerman, K.; Otto, P.; Vidugiris, G.; Zhu, J.; Darzins, A.; Klaubert, D. H.; Bulleit, R. F.; Wood, K. V., HaloTag: a novel protein labeling technology for cell imaging and protein analysis. ACS Chem Biol 2008, 3 (6), 373–82.

30. Pan, C. R.; Knutson, S. D.; Huth, S. W.; MacMillan, D. W. C., microMap proximity labeling in living cells reveals stress granule disassembly mechanisms. Nat Chem Biol 2024.

31. Becker, A. P.; Biletch, E.; Kennelly, J. P.; Julio, A. R.; Villaneuva, M.; Nagari, R. T.; Turner, D. W.; Burton, N. R.; Fukuta, T.; Cui, L.; Xiao, X.; Hong, S. G.; Mack, J. J.; Tontonoz, P.; Backus, K. M., Lipid- and protein-directed photosensitizer proximity labeling captures the cholesterol interactome. bioRxiv 2024.

32. Grimm, J. B.; English, B. P.; Choi, H.; Muthusamy, A. K.; Mehl, B. P.; Dong, P.; Brown, T. A.; Lippincott-Schwartz, J.; Liu, Z.; Lionnet, T.; Lavis, L. D., Bright photoactivatable fluorophores for single-molecule imaging. Nat Methods 2016, 13 (12), 985–988.

33. Zhang, Y.; Wolf-Yadlin, A.; Ross, P. L.; Pappin, D. J.; Rush, J.; Lauffenburger, D. A.; White, F. M., Time-resolved mass spectrometry of tyrosine phosphorylation sites in the epidermal growth factor receptor signaling network reveals dynamic modules. Mol Cell Proteomics 2005, 4 (9), 1240–50.

34. Fraser, J.; Simpson, J.; Fontana, R.; Kishi-Itakura, C.; Ktistakis, N. T.; Gammoh, N., Targeting of early endosomes by autophagy facilitates EGFR recycling and signalling. EMBO Rep 2019, 20 (10), e47734.

35. Palmieri, D.; Bouadis, A.; Ronchetti, R.; Merino, M. J.; Steeg, P. S., Rab11a differentially modulates epidermal growth factor-induced proliferation and motility in immortal breast cells. Breast Cancer Res Treat 2006, 100 (2), 127–37.

36. van Bergen En Henegouwen, P. M., Eps15: a multifunctional adaptor protein regulating intracellular trafficking. Cell Commun Signal 2009, 7, 24.

37. Garvalov, B. K.; Foss, F.; Henze, A. T.; Bethani, I.; Graf-Hochst, S.; Singh, D.; Filatova, A.; Dopeso, H.; Seidel, S.; Damm, M.; Acker-Palmer, A.; Acker, T., PHD3 regulates EGFR internalization and signalling in tumours. Nat Commun 2014, 5, 5577.

38. Guo, H.; Wang, J.; Ren, S.; Zheng, L. F.; Zhuang, Y. X.; Li, D. L.; Sun, H. H.; Liu, L. Y.; Xie, C.; Wu, Y. Y.; Wang, H. R.; Deng, X.; Li, P.; Zhao, T. J., Targeting EGFR-dependent tumors by disrupting an ARF6-mediated sorting system. Nat Commun 2022, 13 (1), 6004.

39. Sousa, L. P.; Lax, I.; Shen, H.; Ferguson, S. M.; De Camilli, P.; Schlessinger, J., Suppression of EGFR endocytosis by dynamin depletion reveals that EGFR signaling occurs primarily at the plasma membrane. Proc Natl Acad Sci U S A 2012, 109 (12), 4419–24.

40. Tice, D. A.; Biscardi, J. S.; Nickles, A. L.; Parsons, S. J., Mechanism of biological synergy between cellular Src and epidermal growth factor receptor. Proc Natl Acad Sci U S A 1999, 96 (4), 1415–20.

41. Wang, T. H.; Lin, Y. H.; Yang, S. C.; Chang, P. C.; Wang, T. C.; Chen, C. Y., Tid1-S regulates the mitochondrial localization of EGFR in non-small cell lung carcinoma. Oncogenesis 2017, 6 (7), e361.

42. Park, O. K.; Schaefer, T. S.; Nathans, D., In vitro activation of Stat3 by epidermal growth factor receptor kinase. Proc Natl Acad Sci U S A 1996, 93 (24), 13704–8.

43. Lo, H. W.; Hsu, S. C.; Ali-Seyed, M.; Gunduz, M.; Xia, W.; Wei, Y.; Bartholomeusz, G.; Shih, J. Y.; Hung, M. C., Nuclear interaction of EGFR and STAT3 in the activation of the iNOS/NO pathway. Cancer Cell 2005, 7 (6), 575–89.

44. Meran, S.; Luo, D. D.; Simpson, R.; Martin, J.; Wells, A.; Steadman, R.; Phillips, A. O., Hyaluronan facilitates transforming growth factor-beta1-dependent proliferation via CD44 and epidermal growth factor receptor interaction. J Biol Chem 2011, 286 (20), 17618–30.

45. Grass, G. D.; Tolliver, L. B.; Bratoeva, M.; Toole, B. P., CD147, CD44, and the epidermal growth factor receptor (EGFR) signaling pathway cooperate to regulate breast epithelial cell invasiveness. J Biol Chem 2013, 288 (36), 26089-26104.

46. Perez, A.; Neskey, D. M.; Wen, J.; Pereira, L.; Reategui, E. P.; Goodwin, W. J.; Carraway, K. L.; Franzmann, E. J., CD44 interacts with EGFR and promotes head and neck squamous cell carcinoma initiation and progression. Oral Oncol 2013, 49 (4), 306–13.

47. Du, W. W.; Fang, L.; Li, M.; Yang, X.; Liang, Y.; Peng, C.; Qian, W.; O’Malley, Y. Q.; Askeland, R. W.; Sugg, S. L.; Qian, J.; Lin, J.; Jiang, Z.; Yee, A. J.; Sefton, M.; Deng, Z.; Shan, S. W.; Wang, C. H.; Yang, B. B., MicroRNA miR-24 enhances tumor invasion and metastasis by targeting PTPN9 and PTPRF to promote EGF signaling. J Cell Sci 2013, 126 (Pt 6), 1440–53.

48. Demosthenous, C.; Han, J. J.; Hu, G.; Stenson, M.; Gupta, M., Loss of function mutations in PTPN6 promote STAT3 deregulation via JAK3 kinase in diffuse large B-cell lymphoma. Oncotarget 2015, 6 (42), 44703–13.

49. Liu, G.; Zhang, Y.; Huang, Y.; Yuan, X.; Cao, Z.; Zhao, Z., PTPN6-EGFR Protein Complex: A Novel Target for Colon Cancer Metastasis. J Oncol 2022, 2022, 7391069.

50. Swaney, D. L.; Ramms, D. J.; Wang, Z.; Park, J.; Goto, Y.; Soucheray, M.; Bhola, N.; Kim, K.; Zheng, F.; Zeng, Y.; McGregor, M.; Herrington, K. A.; O’Keefe, R.; Jin, N.; VanLandingham, N. K.; Foussard, H.; Von Dollen, J.; Bouhaddou, M.; Jimenez-Morales, D.; Obernier, K.; Kreisberg, J. F.; Kim, M.; Johnson, D. E.; Jura, N.; Grandis, J. R.; Gutkind, J. S.; Ideker, T.; Krogan, N. J., A protein network map of head and neck cancer reveals PIK3CA mutant drug sensitivity. Science 2021, 374 (6563), eabf2911.

51. Zhang, Z.; Rohweder, P. J.; Ongpipattanakul, C.; Basu, K.; Bohn, M. F.; Dugan, E. J.; Steri, V.; Hann, B.; Shokat, K. M.; Craik, C. S., A covalent inhibitor of K-Ras(G12C) induces MHC class I presentation of haptenated peptide neoepitopes targetable by immunotherapy. Cancer Cell 2022, 40 (9), 1060–1069 e7.

52. Kiyatkin, A.; van Alderwerelt van Rosenburgh, I. K.; Klein, D. E.; Lemmon, M. A., Kinetics of receptor tyrosine kinase activation define ERK signaling dynamics. Sci Signal 2020, 13 (645).

53. Gerritsen, J. S.; Faraguna, J. S.; Bonavia, R.; Furnari, F. B.; White, F. M., Predictive data-driven modeling of C-terminal tyrosine function in the EGFR signaling network. Life Sci Alliance 2023, 6 (8).

54. Baumdick, M.; Bruggemann, Y.; Schmick, M.; Xouri, G.; Sabet, O.; Davis, L.; Chin, J. W.; Bastiaens, P. I., EGF-dependent re-routing of vesicular recycling switches spontaneous phosphorylation suppression to EGFR signaling. Elife 2015, 4.

55. Ji, H.; Wang, J.; Nika, H.; Hawke, D.; Keezer, S.; Ge, Q.; Fang, B.; Fang, X.; Fang, D.; Litchfield, D. W.; Aldape, K.; Lu, Z., EGF-induced ERK activation promotes CK2-mediated disassociation of alpha-Catenin from beta-Catenin and transactivation of beta-Catenin. Mol Cell 2009, 36 (4), 547–59.

56. Hu, T.; Li, C., Convergence between Wnt-beta-catenin and EGFR signaling in cancer. Mol Cancer 2010, 9, 236.

57. Major, M. B.; Roberts, B. S.; Berndt, J. D.; Marine, S.; Anastas, J.; Chung, N.; Ferrer, M.; Yi, X.; Stoick-Cooper, C. L.; von Haller, P. D.; Kategaya, L.; Chien, A.; Angers, S.; MacCoss, M.; Cleary, M. A.; Arthur, W. T.; Moon, R. T., New regulators of Wnt/beta-catenin signaling revealed by integrative molecular screening. Sci Signal 2008, 1 (45), ra12.

58. Liu, Z.; Ma, L.; Sun, Y.; Yu, W.; Wang, X., Targeting STAT3 signaling overcomes gefitinib resistance in non-small cell lung cancer. Cell Death Dis 2021, 12 (6), 561.

59. Sato, S.; Morita, K.; Nakamura, H., Regulation of target protein knockdown and labeling using ligand-directed Ru(bpy)3 photocatalyst. Bioconjug Chem 2015, 26 (2), 250–6.

60. Muller, M.; Grabnitz, F.; Barandun, N.; Shen, Y.; Wendt, F.; Steiner, S. N.; Severin, Y.; Vetterli, S. U.; Mondal, M.; Prudent, J. R.; Hofmann, R.; van Oostrum, M.; Sarott, R. C.; Nesvizhskii, A. I.; Carreira, E. M.; Bode, J. W.; Snijder, B.; Robinson, J. A.; Loessner, M. J.; Oxenius, A.; Wollscheid, B., Light-mediated discovery of surfaceome nanoscale organization and intercellular receptor interaction networks. Nat Commun 2021, 12 (1), 7036.

61. Trowbridge, A. D.; Seath, C. P.; Rodriguez-Rivera, F. P.; Li, B. X.; Dul, B. E.; Schwaid, A. G.; Buksh, B. F.; Geri, J. B.; Oakley, J. V.; Fadeyi, O. O.; Oslund, R. C.; Ryu, K. A.; White, C.; Reyes-Robles, T.; Tawa, P.; Parker, D. L., Jr.; MacMillan, D. W. C., Small molecule photocatalysis enables drug target identification via energy transfer. Proc Natl Acad Sci U S A 2022, 119 (34), e2208077119.

62. Seath, C. P.; Burton, A. J.; Sun, X.; Lee, G.; Kleiner, R. E.; MacMillan, D. W. C.; Muir, T. W., Tracking chromatin state changes using nanoscale photo-proximity labelling. Nature 2023, 616 (7957), 574–580.

63. Huth, S. W.; Oakley, J. V.; Seath, C. P.; Geri, J. B.; Trowbridge, A. D.; Parker, D. L., Jr.; Rodriguez-Rivera, F. P.; Schwaid, A. G.; Ramil, C.; Ryu, K. A.; White, C. H.; Fadeyi, O. O.; Oslund, R. C.; MacMillan, D. W. C., muMap Photoproximity Labeling Enables Small Molecule Binding Site Mapping. J Am Chem Soc 2023, 145 (30), 16289–16296.

64. Ngo, W.; Peukes, J. T.; Baldwin, A.; Xue, Z. W.; Hwang, S.; Stickels, R. R.; Lin, Z.; Satpathy, A. T.; Wells, J. A.; Schekman, R.; Nogales, E.; Doudna, J. A., Mechanism-guided engineering of a minimal biological particle for genome editing. bioRxiv 2024.

65. da Cunha Santos, G.; Shepherd, F. A.; Tsao, M. S., EGFR mutations and lung cancer. Annu Rev Pathol 2011, 6, 49–69.

66. Rotow, J.; Bivona, T. G., Understanding and targeting resistance mechanisms in NSCLC. Nat Rev Cancer 2017, 17 (11), 637–658.

67. Hrustanovic, G.; Lee, B. J.; Bivona, T. G., Mechanisms of resistance to EGFR targeted therapies. Cancer Biol Ther 2013, 14 (4), 304–14.

68. Haderk, F.; Chou, Y. T.; Cech, L.; Fernandez-Mendez, C.; Yu, J.; Olivas, V.; Meraz, I. M.; Barbosa Rabago, D.; Kerr, D. L.; Gomez, C.; Allegakoen, D. V.; Guan, J.; Shah, K. N.; Herrington, K. A.; Gbenedio, O. M.; Nanjo, S.; Majidi, M.; Tamaki, W.; Pourmoghadam, Y. K.; Rotow, J. K.; McCoach, C. E.; Riess, J. W.; Gutkind, J. S.; Tang, T. T.; Post, L.; Huang, B.; Santisteban, P.; Goodarzi, H.; Bandyopadhyay, S.; Kuo, C. J.; Roose, J. P.; Wu, W.; Blakely, C. M.; Roth, J. A.; Bivona, T. G., Focal adhesion kinase-YAP signaling axis drives drug-tolerant persister cells and residual disease in lung cancer. Nat Commun 2024, 15 (1), 3741.

69. Qiu, S.; Zhao, Z.; Wu, M.; Xue, Q.; Yang, Y.; Ouyang, S.; Li, W.; Zhong, L.; Wang, W.; Yang, R.; Wu, P.; Li, J. P., Use of intercellular proximity labeling to quantify and decipher cell-cell interactions directed by diversified molecular pairs. Sci Adv 2022, 8 (51), eadd2337.

70. Chudnovskiy, A.; Castro, T. B. R.; Nakandakari-Higa, S.; Cui, A.; Lin, C. H.; Sade-Feldman, M.; Phillips, B. K.; Pae, J.; Mesin, L.; Bortolatto, J.; Schweitzer, L. D.; Pasqual, G.; Lu, L. F.; Hacohen, N.; Victora, G. D., Proximity-dependent labeling identifies dendritic cells that drive the tumor-specific CD4(+) T cell response. Sci Immunol 2024, 9 (100), eadq8843.

71. Dai, X.; Liu, Z.; Zhang, S., Over-expression of EPS15 is a favorable prognostic factor in breast cancer. Mol Biosyst 2015, 11 (11), 2978–85.

72. Tian, X.; Yang, C.; Yang, L.; Sun, Q.; Liu, N., PTPRF as a novel tumor suppressor through deactivation of ERK1/2 signaling in gastric adenocarcinoma. Onco Targets Ther 2018, 11, 7795–7803.

73. Liu, X.; He, B.; Xu, T.; Pan, Y.; Hu, X.; Chen, X.; Wang, S., MiR-490-3p Functions As a Tumor Suppressor by Inhibiting Oncogene VDAC1 Expression in Colorectal Cancer. J Cancer 2018, 9 (7), 1218–1230.

74. Geng, Q.; Xian, R.; Yu, Y.; Chen, F.; Li, R., SHP-1 acts as a tumor suppressor by interacting with EGFR and predicts the prognosis of human breast cancer. Cancer Biol Med 2021, 19 (4), 468–85.

75. Irby, R. B.; Yeatman, T. J., Role of Src expression and activation in human cancer. Oncogene 2000, 19 (49), 5636–42.

76. Agoulnik, I. U.; Vaid, A.; Bingman, W. E., 3rd; Erdeme, H.; Frolov, A.; Smith, C. L.; Ayala, G.; Ittmann, M. M.; Weigel, N. L., Role of SRC-1 in the promotion of prostate cancer cell growth and tumor progression. Cancer Res 2005, 65 (17), 7959-67.

77. Ho, J. R.; Chapeaublanc, E.; Kirkwood, L.; Nicolle, R.; Benhamou, S.; Lebret, T.; Allory, Y.; Southgate, J.; Radvanyi, F.; Goud, B., Deregulation of Rab and Rab effector genes in bladder cancer. PLoS One 2012, 7 (6), e39469.

