## Supplemental Information for "Spatiotemporal photocatalytic proximity labeling proteomics reveals ligand-activated extracellular and intracellular EGFR neighborhoods"

### Index

**Supplementary Figures:**
Figure S1. Design of eMultiMap allowed for selective extracellular labeling.
Figure S2. HaloTag-enabled intracellular proximity labeling demonstrated high selectivity in cells.
Figure S3. Rapid interactome changes were captured by eMultiMap.
Figure S4. eMultiMap can capture dynamic EGFR interactomes at different time-points.
Figure S5. Time-resolved iMultiMap interactome profiling facilitated by HaloTagged EGFR.
Figure S6. Genetic manipulation of target genes revealed EGFR ligand-dependent functional neighbors.

**Methods:**
General methods and instrumentation.
Western blot protocol.
General chemical methods and instrumentation.
Antibodies and biological reagents.
Plasmid construction and cell expression.
Cell culture and transfection.
General flow cytometry.
Flag-tagged labeling assay using anti-Flag antibody.
HaloTagged labeling assay using EY-HTL ligand.
Biotinylated protein enrichment.
Sample preparation for LC-MS/MS analysis.
Proteomics analysis of digested peptide samples.
Analysis of proteomics dataset.
EGFR localization analysis.
Proximity ligation assay.
RNA isolation and qRT-PCR experiment.
Software.
Statistical analysis.

**Synthetic Procedures:**
Scheme 1: Synthetic route of EY-HaloTag Ligand (EY-HTL).

**Figure S1: Design of eMultiMap allowed for selective extracellular labeling. (A)** Designs of EGFR constructs used in this study. **(B)** Engineered EGFR constructs, Flag-EGFR and EGFR-HaloTag, were expressed successfully and displayed on the cell surface of A549 cells. Both Flag-EGFR and EGFR-HaloTag contain ecto-displayed Flag-tags for flow cytometry staining. **(C)** General workflow of eMultiMap using an anti-Flag-EY. **(D)** Volcano plots of EGFR neighborhoods using eMultiMap. Labeled proteins in cells expressing Flag-EGFR using anti-Flag-EY were compared with those enriched in WT A549 cells using aryl-diazirine-biotin, aryl-azide-biotin, phenol-biotin, respectively. Significantly enriched proteins are highlighted in red [log2(fold enrichment) ≥ 1, $P < 0.05$, unique peptide ≥ 2, n = 3] and listed in **Table S1-3**.

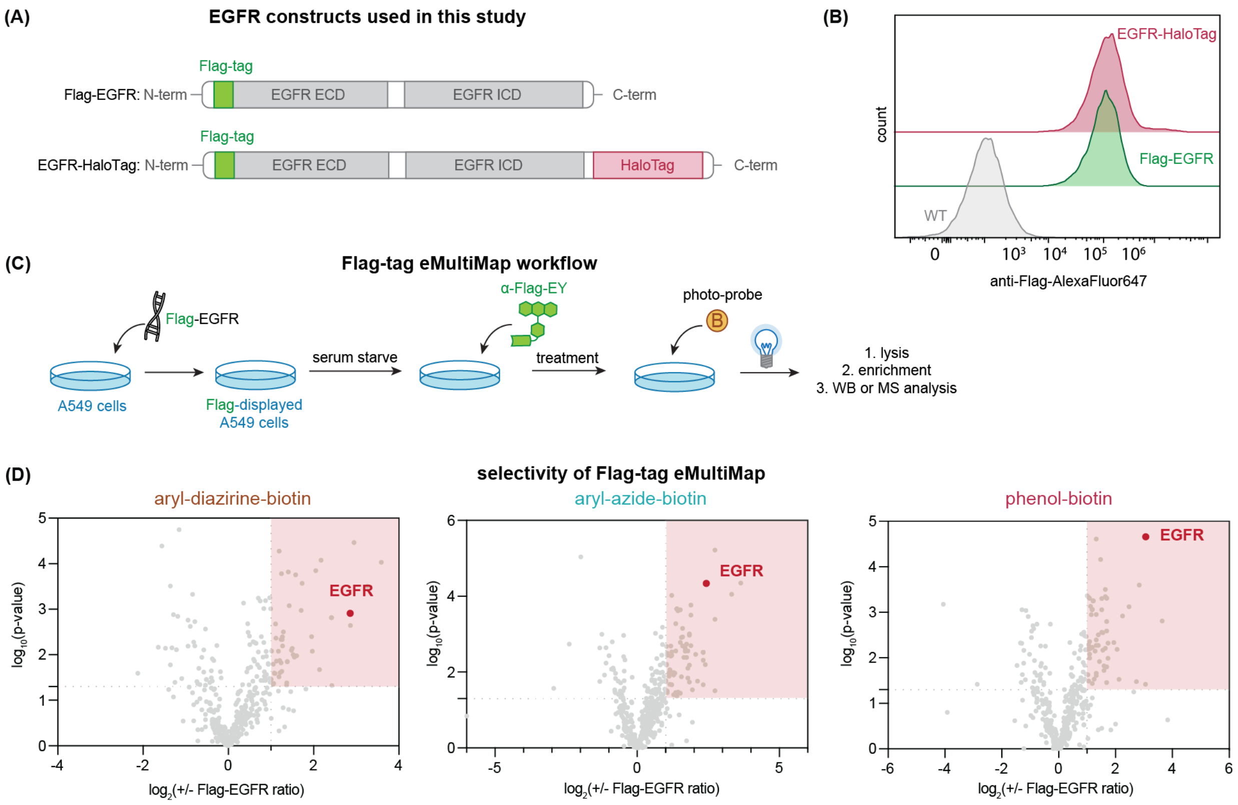

**Figure S2: HaloTag-enabled intracellular proximity labeling demonstrated high selectivity in cells. (A)** Chemical structure of EY-HTL and design of iMultiMap workflow. **(B)** Selective incorporation of HaloTag ligand (HTL) compounds were achieved by repeated wash, demonstrated through differentiation between cells with and without HaloTag expressed. **(C)** General workflow of iMultiMap validation using a GFP-HaloTag construct. Cells expressing either GFP, or EGFP-HaloTag were incubated with EY-HTL before repeated wash and photocatalytic proximity labeling proteomics analysis. **(D)** Volcano plots of EY-HTL-enriched proteins in cells expressing GFP-HaloTag compared to those in cells expressing GFP without HaloTag using aryl-diazirine-biotin, aryl-azide-biotin, and phenol-biotin, respectively. Significantly enriched proteins are highlighted in red [log2(fold enrichment) ≥ 1, P < 0.05, unique peptide ≥ 2, n = 3] and listed in **Table S4-6**.

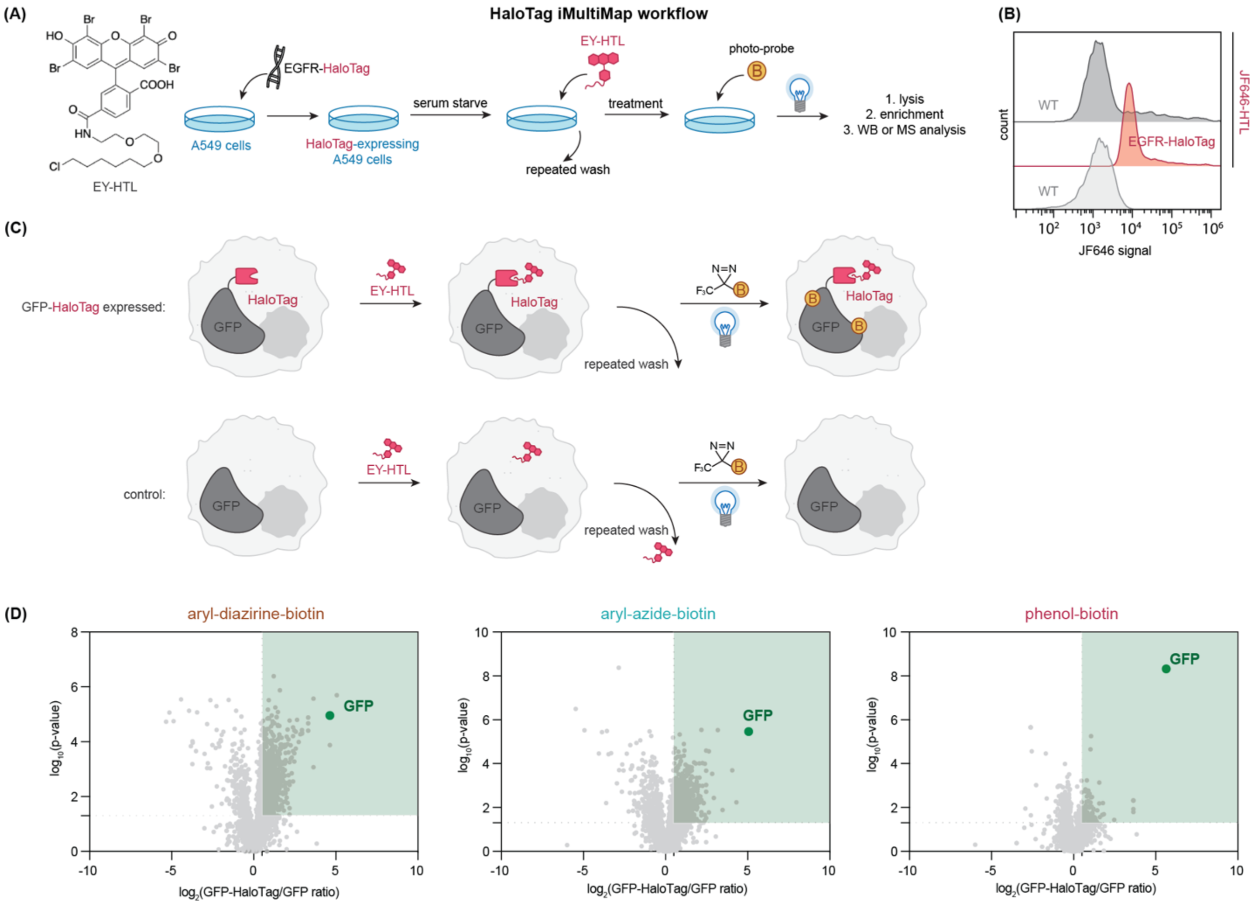

**Figure S3: Rapid interactome changes were captured by eMultiMap. (A)** Localization analysis of Flag-EGFR and EGFR-HaloTag in A549 cells with and without EGF treatment in **Figure 2B**, demonstrating that both constructs are highly co-localized with endogenous EGFR before and after ligand activation. **(B)** Heatmap of the proteins identified in **Figure 2D**. Color indicates enrichment ratio of 5 min EGFR neighborhood over ligand-free conditions, with red indicating increased enrichment and blue indicating decreased enrichment. **(C)** Venn diagram of enriched EGFR interactome with 5 min incubation of EGF enriched from A549 cells by using different photo-probes.

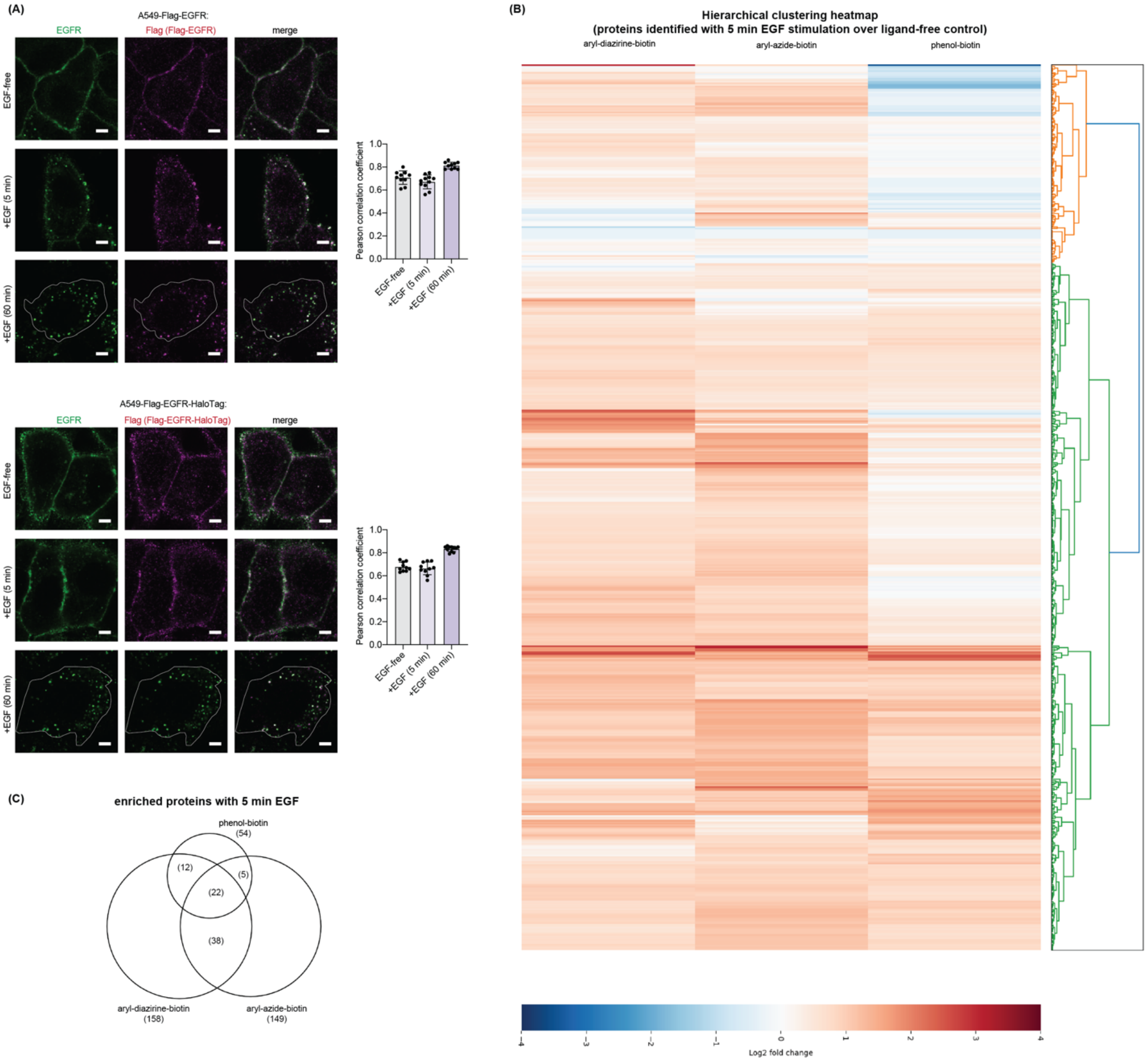

**Figure S4: eMultiMap can capture dynamic EGFR interactomes at different time-points. (A)** Heatmap of the proteins identified in **Figure 3A** compared to that from **Figure 2D**. Color indicates enrichment ratio of treatment over ligand-free conditions, with red indicating increased enrichment and blue indicating decreased enrichment. **(B)** Venn diagram of enriched EGFR interactome with 60 min incubation of EGF enriched from A549 cells by using different photo-probes. **(C)** Direct comparison of enriched EGFR interactome captured with 5 min EGF and that with 60 min EGF. **(D)** Representative images of proximity ligation assay visualizing proximity of EGFR and EGFR targets, Rab11a, EPS15, PTPRF, STAT3 and Src shown in **Figure 3C**, respectively. PLA puncta are in green, indicating proximity between target and EGFR, and nuclei are in blue. Scale bar = 20 μm.

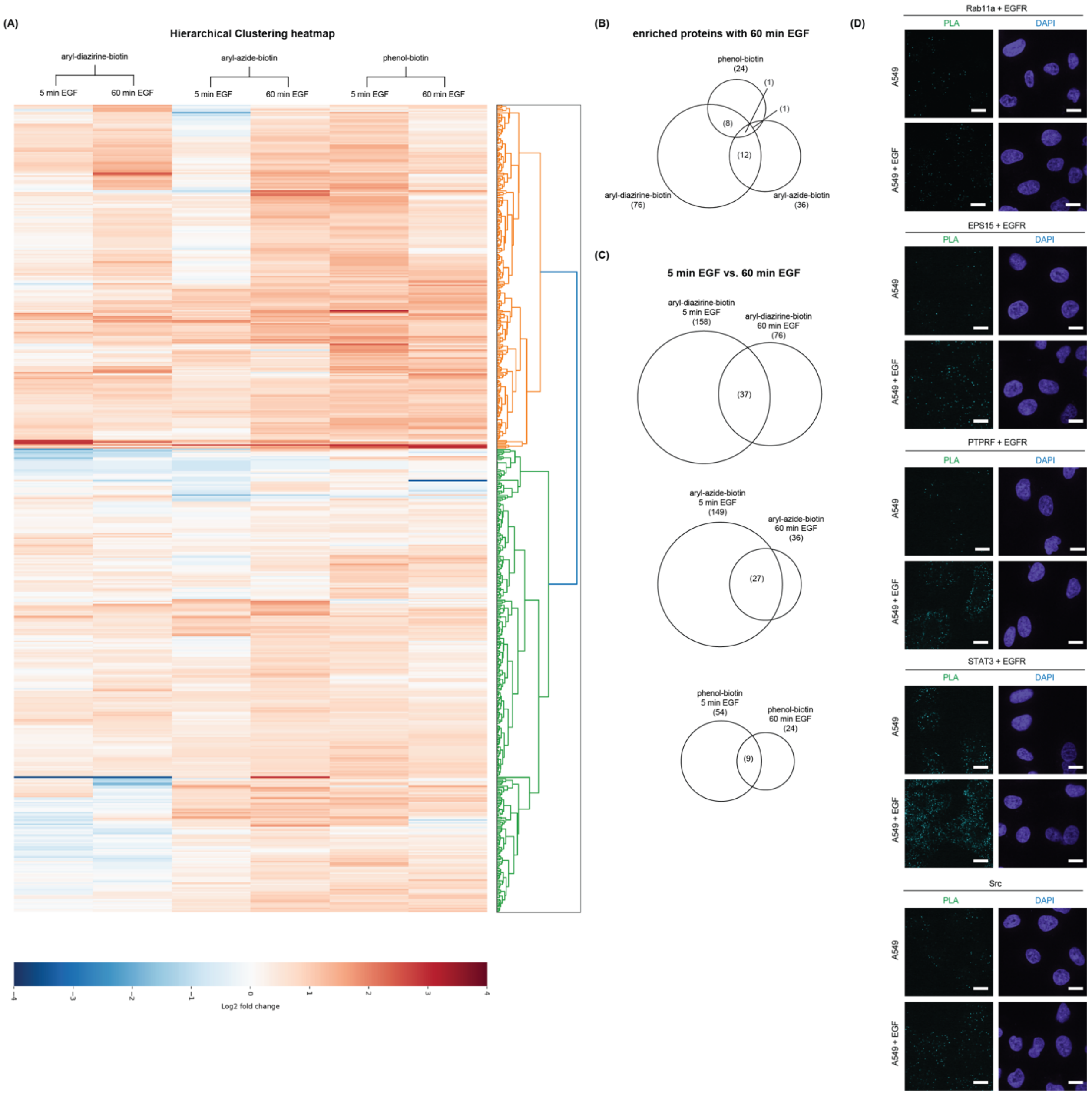

**Figure S5: Time-resolved iMultiMap interactome profiling facilitated by HaloTagged EGFR. (A)** We integrated the time-resolved ligand treatment workflow with iMultiMap design, which enabled highly controlled photocatalytic proximity labeling in cells. **(B)** Volcano plots of rapid changes of on-state EGFR neighborhoods using iMultiMap. Labeled proteins using EY-HTL were compared with or without 5 min EGF stimulation using aryl-diazirine-biotin, aryl-azide-biotin, phenol-biotin, respectively. Significantly enriched proteins are highlighted in red [log2(fold enrichment) ≥ 1, P < 0.05, unique peptide ≥ 2, n = 3] and listed in **Table S16-18**. **(C)** Volcano plots of prolonged changes of on-state EGFR neighborhoods using eMultiMap. Labeled proteins using anti-Flag-EY were compared with or without 60 min EGF stimulation using aryl-diazirine-biotin, aryl-azide-biotin, phenol-biotin, respectively. Significantly enriched proteins are highlighted in red [log2(fold enrichment) ≥ 1, P < 0.05, unique peptide ≥ 2, n = 3] and listed in **Table S19-21**.

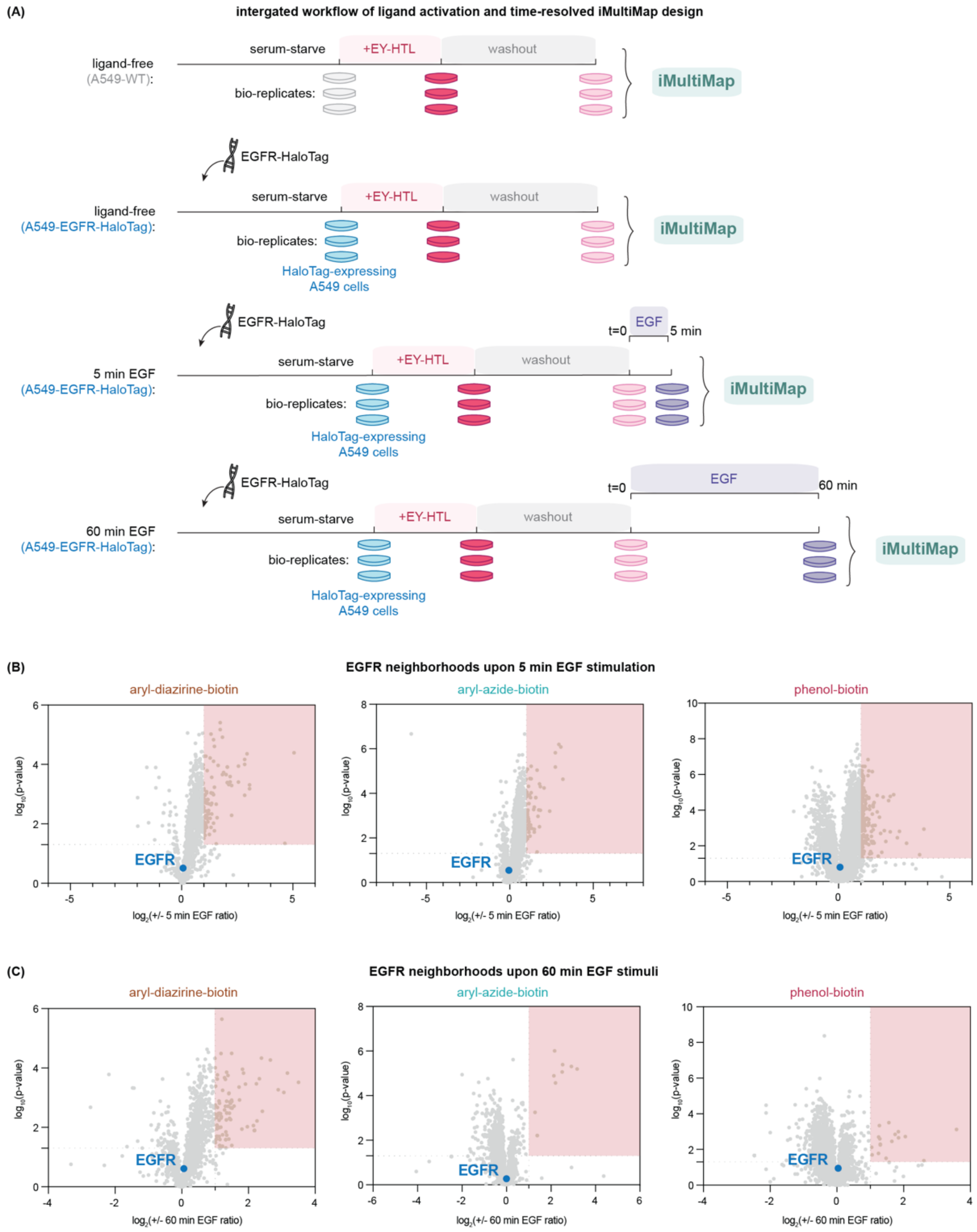

**Figure S6: Genetic manipulation of target genes revealed EGFR ligand-dependent functional neighbors. (A)** WB validation of knock-down of SRC, PTPRF or ARF6 in A549 cells used in **Figure 5B**. **(B)** Establishment of a time-course flow cytometry workflow visualizing cell surface EGFR in WT A549 cells. **(C)** WB validation of knock-down of EPS15, Rab11a or DYN1 in A549 cells used in **Figure 5C**. **(D)** Representative flow cytometry of EGFR internalization assay monitoring cell surface EGFR upon EGF stimulation in cells with EPS15, Rab11a, DYN1 or ARF6 knocked down. **(E)** Changes of EGFR level upon EGF stimulation and cycloheximide (100 ng/µL) to monitor EGFR degradation in cells with EPS15, Rab11a, DYN1 or ARF6 knocked down. **(F)** WB validation of EGFR knock-down used in **Figure 5D**.

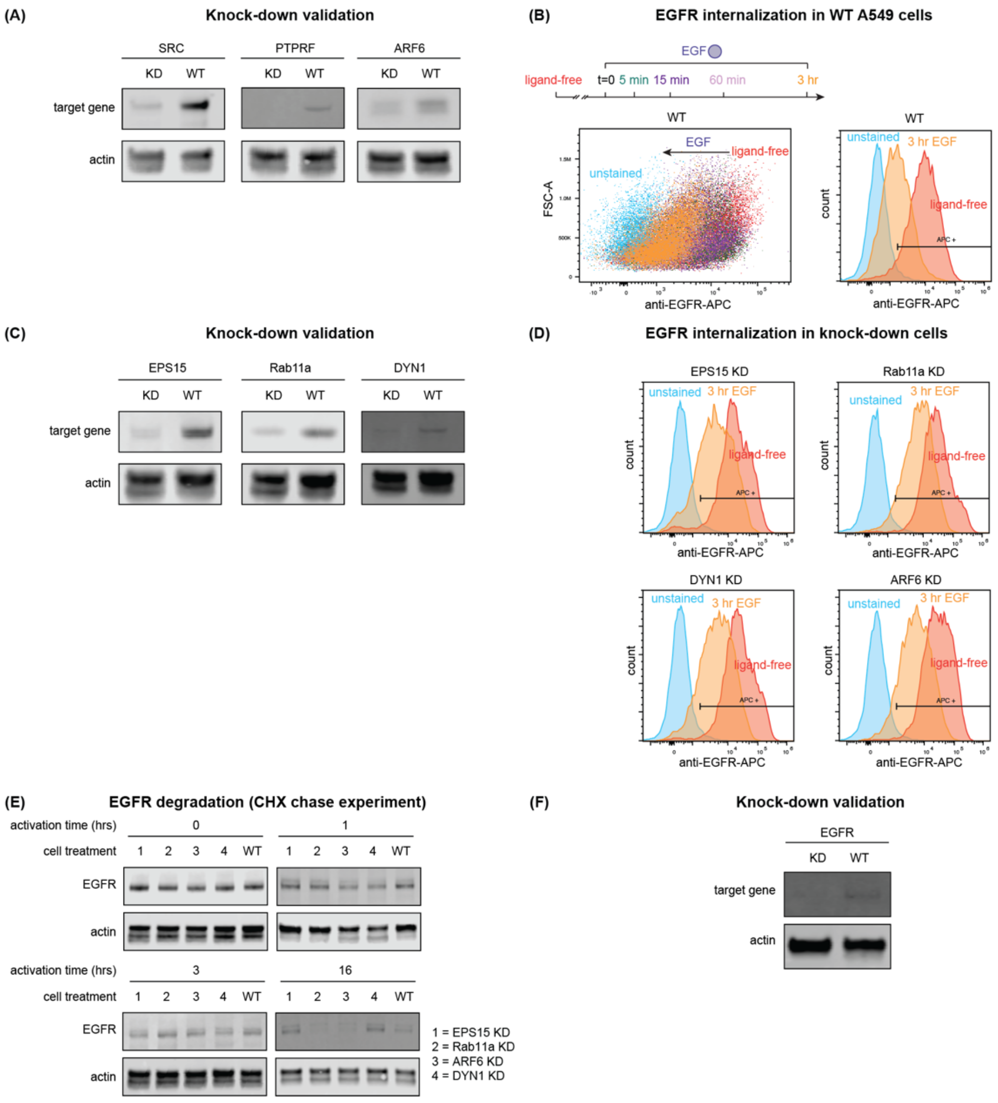

#### Methods:

##### General methods and instrumentation.

Illumination was performed using a Penn PhD Photoreactor M2 (Sigma Aldrich, Z744035) with a 450 nm blue light source module (Sigma Aldrich, Z744033) at 100% intensity, or LED array light sources (Thor Labs, LIU470A for 470 nm LED array, LIU525B for 525 nm LED array) along with a LED mounting adapter (AD38). To use the Penn PhD Photoreactor, fan speed was set at 6800 rpm under manual control with 100/min stirring and samples were illuminated at 100% intensity for indicated time. To use LED array light sources, samples were placed under a Thor Labs LED array light source, which provides 4.0 $mW/cm^2$ (470 nm) and 1.9 $mW/cm^2$ (525 nm) intensity at 100 mm distance from the LED according to information from the manufacturer. Flow cytometry experiments were performed on a CytoFlex flow cytometer (Beckman CytoFlex) and analyzed using FlowJo software. Proteomics experiments were performed on a TimsTOF PRO (Bruker) equipped with a CaptiveSpray source and a nanoElute System. The peptides were separated on a 25 cm, ReproSil c18 1.5 µM 100 Å column (PepSep, PN. # PSC- 25-150-15-UHP-nc). Protein quantification was performed by bicinchoninic acid assay on a multimode microplate reader Infinite 200 PRO (Tecan Trading AG, Switzerland). Sonication of cells or protein pellets was performed using a QSonica Q500 Sonicator (QSonica Sonicators, Newtown, CT). DNA or protein concentrations were measured using a NanoDrop 2000 spectrophotometer (Thermo Scientific).

##### Western blot protocol.

For immunoblotting analysis, proteins were loaded on 4-12% 17-well BisTris gels (Thermo Fischer, NW04127BOX), and transferred from SDS-PAGE gels to PVDF membranes (Thermo Fischer, IB24002) using an iBlot-2 dry blotting system (Thermo Scientific, IB21001). Membranes were blocked with Intercept® (TBS) blocking buffer (LI-COR, 927-60001), incubated with the primary antibodies as indicated by vendors, washed with Tris buffered saline containing 0.1% Tween-20 (TBST, 37mM sodium chloride, 20mM Tris, 2.7mM potassium chloride, 0.05% Tween 20; pH=7.4) and incubated with secondary antibodies sequentially including anti-rabbit IgG Goat IR800 secondary antibody (Rockland, 926-32211), anti-rabbit IgG Goat IR680 secondary antibody (Rockland, 611-144-002), anti-rabbit IgG Goat secondary antibody peroxidase (Rockland, 611-1302), anti-mouse IgG Goat IR800 secondary antibody (Rockland, 610-145-211) and/or anti-mouse IgG Goat IR680 secondary antibody (Rockland, 610-144-002). Immunoblots images were captured by an infrared LI-COR imager (Odyssey CLx). In-gel fluorescence and immunoblot fluorescence signals were detected on a BioRad imager (ChemiDoc XRS+ System). Analysis was performed using ImageStudioLite.

##### General chemical methods and instrumentation.

Chemicals were purchased including TFPA-$PEG_3$-biotin (Thermo Scientific, 21303), biotinyl tyramide (Sigma-Aldrich, SML2135) and Eosin Y (Sigma-Aldrich, E4009). DBCO-$PEG_3$-EY was synthesized and characterized by ChemPartner as previously reported. Aryl-diazirine-biotin (diazirine-$PEG_3$-biotin) was synthesized and characterized by Medicilon as previously reported. Eosin Y-HaloTag ligand was synthesized in house first before being batch synthesized and characterized by ChemPartner as shown in Scheme 1. All solvents and reagents were purchased from chemical suppliers (Sigma Aldrich, Acros Organics, Thermo Scientific or VWR Chemicals BDH®) and were used as received unless otherwise noted. Flash Column Chromatography was performed using Teledyne ISCO CombiFlash EZ Prep chromatography system, employing pre-packed silica gel Teledyne ISCO RediSep cartridges. Protein mass spectra were obtained using a Waters Xevo G2-XS time-of-flight mass spectrometer operating with Waters MassLynx software (version 4.2).

##### Antibodies and biological reagents.

Antibodies were purchased including: EGFR (Thermo Scientific, MA5-13319; Cell Signaling Technology, 4267S, 2239S, 2256S), Rab11a (Cell Signaling Technology, 2413S), STAT2 (Cell Signaling Technology, 72604T), EPS15 (Cell Signaling Technology, 12460S), SHP1 (Cell Signaling Technology, 26516S), ITGA2 (Bethyl Laboratories, A305-211A-T), Dynamin I/II (Cell Signaling Technology, 2342S), SRC (Cell Signaling Technology, 2109S), ARF6 (Cell Signaling Technology, 5740S), PTPRF (Cell Signaling Technology, 61611S), β-actin (Santa Cruz Biotechnology, sc-47778), CD44 (Cell Signaling Technology, 3578S), Tid1 (Cell Signaling Technology, 4775S), β-catenin (Cell Signaling Technology, 8480T), GRB2 (Cell Signaling Technology, 3972S), Phospho-EGFR (Tyr1045) (Cell Signaling Technology, 2237T), Phospho-EGFR (Tyr1086) (Cell Signaling Technology, 2220S), Phospho-EGFR (Tyr845) (Cell Signaling Technology, 2231S), Phospho-EGFR (Tyr992) (Cell Signaling Technology 2235T), Phospho-EGFR (Tyr1068) (Cell Signaling Technology, 3777T), Phospho-EGFR (Tyr992) (Cell Signaling Technology, 2235T), Phospho-EGFR (Tyr978) (Cell Signaling Technology, 3790), Phospho-EGFR (Tyr1173) (Cell Signaling Technology, 4757), Phospho-p44/42 MAPK (Erk1/2) (Thr202/Tyr204) (Cell Signaling Technology, 4370S), P44/42 MAPK (Erk1/2) (Cell Signaling Technology, 4695T), Phospho-EGFR (Y1068) Fluorescein-conjugated Antibody (R&D Systems, IC3570F), Phospho-EGFR Pathway Antibody panel (Cell Signaling Technology, 9789T). Anti-M1-FLAG monoclonal antibody was purified from M1 hybridoma as previously reported.

The following antibodies were used in flow cytometry assays: streptavidin-AlexaFluor488 (Thermo Scientific, S32354), streptavidin-AlexaFluor647 (BioLegend, 405237), anti-EGFR-AlexaFluor647 (Fisher Scientific, 352918), anti-Flag-AlexaFluor647 (BioLegend, 637316). Other biological reagents such as recombinant Human EGF Protein (R&D Systems, 236-EG-200), Cetuximab (Selleck Chemicals, A2000), fluorescein-conjugated EGF (Thermo Scientific, E3478), Cycloheximide (Sigma-Aldrich, 01810-1G), flow cytometry fixation buffer (R&D Systems, FC004), Triton X-100 (1%) (Thermo Scientific, HFH10), Pierce™ High Capacity Streptavidin Agarose (Thermo Scientific, 20359), FITC Conjugation Kit (Abcam, ab188285), AlexaFluor 647 Conjugation Kit (Abcam, ab269823), Opti-MEM™ I Reduced Serum Medium (Thermo Scientific 31985062), TransIT-PRO® Transfection Reagent (Mirus Bio, MIR-5740), Lipofectamine RNAiMAX Transfection Reagent (Thermo Scientific, 13778075) were purchased. Janelia Fluor-646 (JF646) with HaloTag ligand was a gift from the Lavis lab at Janelia.

For enrichment assays, Pierce™ Streptavidin Agarose beads (Pierce, 20349) and mini Bio-Spin columns (Bio-Rad, 732-6207) were used. Cell lysis buffer was prepared by diluting from 10X cell lysis buffer (Cell Signaling Technology, 9803S) or from 10X RIPA buffer (EMD Millipore, 20-188) and supplemented with protease inhibitor (Protease Inhibitor Cocktail 100X, Cell Signaling Technology, 5871) or phosphatase inhibitor (Phosphatase Inhibitor Cocktail 100X, Cell Signaling Technology, 5870) as indicated. Sample loading buffer was diluted from 4X Laemmli sample loading buffer (Bio-Rad, 161-0747). When performing solvent exchange processes, 7 kDa Zeba Spin desalting columns (ThermoFisher, 89883) were used.

**Plasmid construction and cell expression.**
Plasmids for the Flag-EGFR and Flag-EGFR-HaloTag were constructed in a pcDNA3.1(+) backbone by standard molecular biology methods using its native signal sequence with a GGGGS link between tags and EGFR sequence.

For gene knockout or knockdown, Cas9 constructs with sgRNA spacer sequences and commercial siRNAs (ThermoFisher Scientific, AM16708) were used as previously reported.[1] In short, spacer sequences listed below were cloned into the pCF822-recipient_U6-sgRNA-EFS-Cas9-P2A-Puro backbone (Addgene # 211645): Spacer sequences and their reverse complements were ordered with 5'CACC and 5'AAAC overhangs, then phosphorylated and annealed by incubating with T4 PNK (NEB M0201) (10 U) in T4 DNA Ligase Buffer (NEB B0202S) (10 µL, 1X). The pCF822-recipient_U6-sgRNA-EFS-Cas9-P2A-Puro backbone was digested with Esp3I (NEB R0734) (10 U) and dephosphorylated using CIP (NEB M0525) (6 U) for 2 hours at 37°C. The backbone was subsequently gel purified (Qiagen 28704). Backbone (50 ng) and phosphorylated spacers (0.5 pmol) were annealed with T4 DNA ligase (M0202) (400 U) for 2 hours at 25°C. The ligated plasmids were transformed into Mach1 competent cells (ThermoFisher Scientific, C862003). Plasmids were verified by whole plasmid sequencing after purification using either mini-prep (Qiagen 27104) or midi-prep kits (Zymo, D4200). All sequences were confirmed by Sanger (Quintarabio) and whole-plasmid (Primordium Labs) sequencing.

| sgRNA spacer sequences | |
|---|---|
| Gene | Spacer sequence |
| SRC | TAGCAACAAGAGCAAGCCCA |
| SRC | GTCATAGAGGGCCACAAAGG |
| PTPRF | GTCGATGGAAGGGAACCCAG |
| PTPRF | GGGCACCATCGTCCTCCCTG |
| EPS15 | TGATGGAAACACCATGGCTG |
| EPS15 | TTAGCCGACACAGATGGCAA |
| RAB11A | TGTTGCAAACTCTACTCCAA |
| RAB11A | CCATGGCCTCACCTTTAAAG |
| EGFR | GATGTTCAATAACTGTGAGG |
| EGFR | AGCGATGCGACCCTCCGGGA |
| ARF6 | GGTGACCACCATTCCCACTG |
| ARF6 | TGTGGGTTTCAACGTGGAGA |
| DNM1 | GGCCAGGTCAGAGTTGGCGG |
| DNM1 | CATGGAAGATCTCATCCCGC |

**Cell culture and transfection.**
A549 cells were acquired from the UCSF cell culture and banking services and cultured in Dulbecco's Modified Eagle Medium (DMEM, Thermo Fisher, 11995073), supplemented with penicillin (50 µg/mL), streptomycin (50 µg/mL), and 10% (v/v) FBS. Transfection was performed using TransIT-PRO (Mirus Bio, fMIR 5740) according to the manufacturer’s instructions.

**General flow cytometry.**
Cultured cells were incubated at 37 °C in 5% $CO_2$ for the duration of the assay. To guarantee minimal cell activity, cells were moved onto ice right after treatment and washed three times with 1 mL pre-chilled PBS. Then pelleted cells were blocked with 1 mL filtered 3% BSA in PBS for 30 minutes at 4 °C and stained with corresponding antibody for 1 h at 4 °C. Stained cells were then washed three times with 1 mL pre-chilled PBS and resuspended in 0.5 mL PBS for flow cytometry analysis. Flow cytometry data were analyzed on FlowJo.

**Flag-tagged labeling assay using anti-Flag antibody.**
Transfected A549 cells were incubated at 37 °C in 5% $CO_2$ to 80% confluency and serum-starved overnight. They were then washed with pre-chilled PBS three times. Indicated amounts of anti-Flag antibody or antibody-EY conjugate were pre-chilled and added to the cells for 15 min at 4 °C before excessive antibody-EY conjugates were removed by washing with pre-chilled PBS. The

antibody-bound cells were then resuspended in serum-free DMEM media and treated as indicated. Cells were then dissociated and collected at 4 °C before indicated photo-probe reagents (aryl-diazirine-biotin, aryl-azide-biotin or biotin-phenol) were added to reach a final concentration of 100 μM and let incubate for 15 min at 4 °C. Thoroughly mixed cells were illuminated with LED for indicated time at 4 °C, pelleted and subjected to flow cytometry or LC-MS/MS sample preparation.

**HaloTagged labeling assay using EY-HTL ligand.**

Transfected A549 cells were incubated at 37 °C in 5% $CO_2$ to 80% confluency and serum-starved overnight. They were then washed with PBS three times and incubated with indicated concentrations of EY-HTL for 30 min at 37 °C. Then the excessive molecules were washed off with serum-free media twice and incubated with serum-free media for 15 min at 37 °C, which was subsequently repeated four times. Cells were then treated as indicated and collected at 4 °C before indicated photo-probe reagents (aryl-diazirine-biotin, aryl-azide-biotin or biotin-phenol) were added to reach a final concentration of 100 μM and incubated for 15 min at 4 °C. Thoroughly mixed cells were illuminated with LED for indicated time at 4 °C, pelleted and subjected to flow cytometry or LC-MS/MS sample preparation.

**Biotinylated protein enrichment**

To enrich biotinylated proteins, cell pellets were resuspended in 1 mL 1X RIPA lysis buffer (EMD Millipore) supplemented with protease inhibitor (Protease Inhibitor Cocktail 100X, Cell Signaling Technology, 5871). After 15 min incubation on ice, cells were sonicated for 15 sec (5 sec on, 5 sec off, 20%). Cell lysates were then cleared by centrifugation at 20,000 g for 10 min at 4 ˚C. Proteins were then added to 150 μL Pierce™ Streptavidin Agarose beads (Pierce, 20349) that were pre-washed with 1 mL PBS for three times and incubated for 16 h at 4 ˚C. Afterwards, supernatant was discarded using mini Bio-spin columns (Bio-Rad) and the beads were washed three times with 1 mL 1X RIPA lysis buffer, three times with 1 mL 1M NaCl in 1X PBS, and three time with 1 mL of freshly prepared 2M urea in 50 mM ammonium bicarbonate. The beads were then subjected to Western blotting analysis or LC-MS/MS analysis.

**Sample preparation for LC-MS/MS analysis.**

Proteins on the washed beads were then digested using the Preomics iST kit in an on-bead digestion format according to the manufacturer's instructions. In brief, washed beads were suspended in 100 μL LYSE buffer provided by the Preomics iST kit and incubated at 55 ˚C for 10 min for reduction and alkylation. Once the beads cooled down to room temperature, 50 μL of pre-reconstituted DIGEST were added to the beads and incubated at 37 ˚C for 3 hours with shaking. The digested peptides were then collected using mini Bio-Spin columns (Bio-Rad) and another 50 μL of LYSE buffer were added to wash the beads. Afterwards, 100 μL of STOP solution was added to the combined flow-through elution and mixed using vigorous vortexing. Then the peptides were desalted using the Preomics desalting columns before they were dried under vacuum and resuspended in 15 μL solvent A (0.1% formic acid with 2% acetonitrile) for mass spectrometry analysis. Peptide amount was monitored using NanoDrop 2000 spectrophotometer (Thermo Scientific).

**Proteomics analysis of digested peptide samples.**

Proteomics experiments were performed on a TimsTOF PRO (Bruker) equipped with a CaptiveSpray source and a nanoElute system. The peptides were separated on a 25 cm, ReproSil c18 1.5 μM 100 Å column (PepSep, PN. # PSC-25-150-15-UHP-nc) using a stepwise linear gradient method with water in 0.1% formic acid (solvent A) and acetonitrile with 0.1% formic acid (solvent B): 5-30% solvent B for 90 min at 0.5 μL/min, 30-35% solvent B for 10 min at 0.6 μl/min, 35-95% solvent B for 4 min at 0.5 μL/min, 95% hold for 4 min at 0.5 μL/min). Acquired data was collected in a data-dependent acquisition mode with ion mobility activated in PASEF mode. MS and MS/MS spectra were collected with m/z ranging from 100 to 1700 in positive mode.

**Analysis of proteomics dataset.**
All acquired data was searched using PEAKS online Xpro 1.6 (Bioinformatics Solutions Inc.) or FragPipe powered by MSFragger (v3.7). Spectral searches were performed using a curated FASTA-formatted dataset containing Swiss Uniprot-reviewed human proteome file with gene ontology localized on the plasma membrane (downloaded from UniProt database). A precursor mass error tolerance was set to 20 ppm and a fragment mass error tolerance was set at 0.03 ppm. Peptides, ranging from 6 to 45 amino acids in length, were searched in semi-specific trypsin digest mode with a maximum of three missed cleavages. Carbamidomethylation (+57.0214 Da) on cysteines was set as a static modification while methionine oxidation (+15.9949 Da) and lysine acetylation (+42.0115 Da) were set as a variable modification. Peptides were filtered based on a false discovery rate (FDR) of 1%. Samples were normalized using total ion current (TIC) or EGFR. For p-value and fold change calculations, the data were further processed using a customized script as previously described.[2]

**EGFR localization analysis**.
EGFR localization analysis was performed using A549 cells plated on an 18-well chamber slide precoated with ibiTreat (iBidi, 81816) that were serum-starved overnight before treatment and with or without EGF for indicated time. Cells were then fixed with 4% paraformaldehyde freshly diluted from 16% paraformaldehyde (Electron Microscopy Sciences, 15710) for 10 min at 25 °C and permeabilized with 0.1% Triton in PBS for 15 min at 25 °C before blocking with 3% goat serum albumin (GSA) for 1 hour at 4 °C. Primary antibodies diluted in 3% GSA were incubated at 4 °C overnight, followed by washes and incubation with diluted secondary antibodies for 1 hour at 4 °C. Cells were finally stained with DAPI before confocal imaging. Confocal microscopy was performed using a 60X Plan-Apo 679 1.42 NA oil objective on an Olympus Fluoview 3000 laser scanning confocal microscope. Images were acquired as Z-stacks using Galvano scanning mode and confocal zoom of 5X magnification. Individual cells were manually segmented, and co-localization was quantified with Pearson correlation coefficient using the Coloc2 plugin in ImageJ.

**Proximity ligation assay.**
Proximity ligation assay was performed according to published literature.[3, 4] In brief, A549 cells were plated on an 18-well chamber slide precoated with ibiTreat (iBidi, 81816) and serum-starved overnight before treatment with or without EGF for indicated time. Cells were then fixed with 4% paraformaldehyde freshly diluted from 16% paraformaldehyde (Electron Microscopy Sciences, 15710) for 10 min at 25 °C and permeabilized with 0.1% Triton in PBS for 15 min at 25 °C before a Duolink in situ PLA assay kit (Sigma-Aldrich, DUO92014-30RXN) was used according to the manufacturer's instructions. In brief, the cells were blocked with 1X Duolink Blocking Buffer for 60 min at 37 °C before primary antibodies diluted with Duolink antibody diluent as indicated by antibody vendors were added. After incubation at 4 °C overnight, cells were washed with Duolink Wash Buffer A (Sigma-Aldrich, DUO82049-4L) and diluted PLA secondary antibodies (anti-mouse MINUS, DUO92004-30RXN; anti-rabbit PLUS, DUO92002-30RXN) were added and incubated for 60 min at 37 °C. After washes with Duolink Wash Buffer A, ligation reaction was performed with supplied reagents from Duolink in situ green detection kit (Sigma-Aldrich, DUO92014-30RXN) for 30 min at 37 °C followed by amplification reaction for 100 min at 37 °C. Cells were then washed with Duolink Wash Buffer B (Sigma-Aldrich, DUO82049-4L) and 100-fold diluted Duolink Wash Buffer B before the samples were stained with Duolink mounting solution with DAPI (Sigma-Aldrich, DUO82040-5ML) for 15 min in dark. Images were obtained on an inverted Nikon Ti-E microscope equipped with a Yokogawa CSU-22 confocal scanner unit. PLA spots in cells were counted automatically using imageJ QuickPALM function of analyzing particles with minimum

SNR set as 20.00 and maximum FWHM set as 7 px on at least 5 individual images. Quantification was demonstrated as average ± SD from an average of 200 cells. Image acquisition parameters and data processing parameters were the same for all treatment conditions.

**RNA isolation and qRT-PCR experiment.**
Total RNA was extracted from cells with indicated treatment using RNeasy Mini Kit (Qiagen, 74106), QIAshredder (Qiagen, 79656) and RNase-Free DNase Set (Qiagen, 79254). Reverse transcription was performed using Superscript II Reverse Transcriptase (Thermo Fisher Scientific, 18064014). qPCR was carried out using the TaqMan fast Advanced Master Mix (Applied Biosystems, 44-445-57) on a QuantStudio 12K Flex Real-Time qPCR system (Applied Biosystems). Target gene expression was measured with specific TaqMan probes for AXIN2 (Hs00610344_m1), ZEB1(Hs01566408_m1), cMYC (Hs00153408_m1), and GAPDH (Hs02786624_g1). Relative gene expression levels were calculated using the ΔΔCt method and normalized to the housekeeping gene GAPDH.

**Software.**
Data was analyzed and visualized using GraphPad Prism (v8.0.1) and Microsoft Excel (v16.22), in addition to software listed by each experiment. NMR data were analyzed using MestReNova (v15.0.0). DNA and protein sequences were analyzed using SnapGene (v6.0.2). Flow cytometry data were analyzed by FlowJo (v10.6.1). Proteomics data were analyzed by PEAKS online (Xpro 1.6) and FragPipe powered by MSFragger (v3.7). Images were made using ImageStudioLite (v5.2.5), Adobe Illustrator (v22.1) and BioRender (v2.0).

**Statistical analysis.**
Statistical analyses (unpaired Student's *t*-tests) were performed using GraphPad Prism. Data were derived from at least three biological replicate experiments and presented as the average ± SD, P≤ 0.05, **P≤ 0.01, ***P ≤ 0.001, ****P ≤ 0.0001 and n.s., not significant.

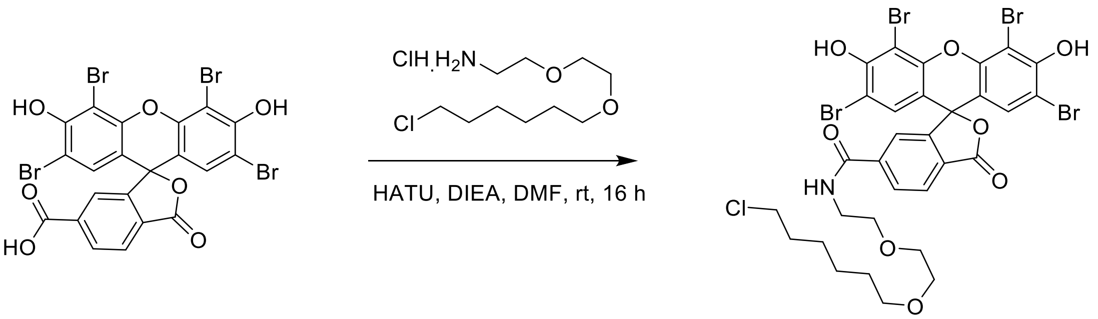


**Scheme 1: Synthetic route of EY-HaloTag Ligand (EY-HTL).**

**Synthetic procedures:**
Synthesis of EY-HaloTag Ligand (EY-HTL), 2',4',5',7'-tetrabromo-N-(2-(2-[(6-chlorohexyl)oxy]ethoxy)ethyl)-3',6'-dihydroxy-3-oxo-3H-spiro[2-benzofuran-1,9'-xanthene]-6-carboxamide.

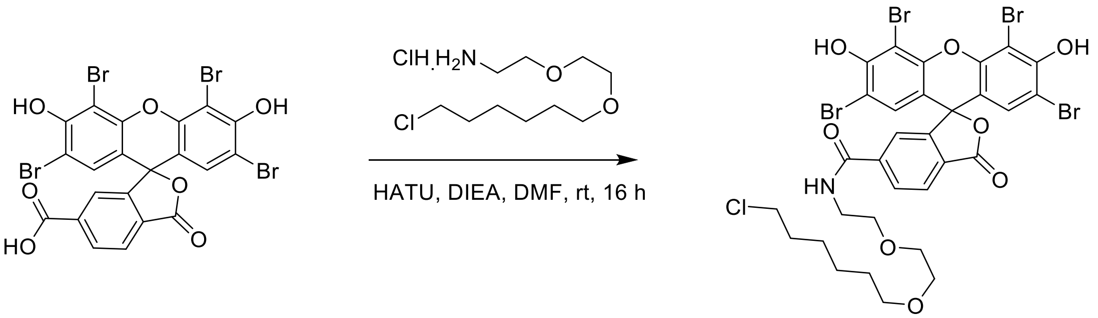


EY-HTL was synthesized similarly to reported literature.[5] To a mixture of 2',4',5',7'-tetrabromo-3',6'-dihydroxy-3-oxo-3H-spiro[2-benzofuran-1,9'-xanthene]-6-carboxylic acid (50mg, 0.072mmol), DIPEA (28.02mg, 0.22mmol) and 1-(2-(2-aminoethoxy)ethoxy)-6-chlorohexane (20.68mg, 0.092mmol) in DMF (5mL) was added HATU (32.97mg, 0.087mmol) in portions at 0 ℃ under Ar atmosphere. The resulting mixture was stirred for 16 hours at room temperature under Ar atmosphere. The residue was purified by C18 column chromatography eluted with ACN: $H_2O$ (5% ~95%) to afford desired product (23mg, yield 35.46%) as a red solid. Synthesis and characterization was performed by Medicilon. LCMS: IDSUF06-WX-MS, Method: 95_15 min, t=6.515 min, MS (ESI): m/z=897.7 $[M+H]^+$. $^1$H NMR (400 MHz, DMSO-$d_6$): δ 10.83 (s, 1H), 8.76 (*d*, *J* = 46.5 Hz, 1H), 8.82-8.71 (m, 1H), 8.24-8.09 (m, 2H), 7.85-7.80 (m, 1H), 7.02 (s, 2H), 3.48-3.35 (m 10H), 3.30-3.27 (m, 2H), 1.67-1.63 (m, 2H), 1.41-1.23 (m, 6H).

**References:**


1. Tan, I. L.; Perez, A. R.; Lew, R. J.; Sun, X.; Baldwin, A.; Zhu, Y. K.; Shah, M. M.; Berger, M. S.; Doudna, J. A.; Fellmann, C., Targeting the non-coding genome and temozolomide signature enables CRISPR-mediated glioma oncolysis. *Cell Rep* **2023,** *42* (11), 113339.
2. Lin, Z.; Schaefer, K.; Lui, I.; Yao, Z.; Fossati, A.; Swaney, D. L.; Palar, A.; Sali, A.; Wells, J. A., Multiscale photocatalytic proximity labeling reveals cell surface neighbors on and between cells. *Science* **2024,** *385* (6706), eadl5763.
3. Swaney, D. L.; Ramms, D. J.; Wang, Z.; Park, J.; Goto, Y.; Soucheray, M.; Bhola, N.; Kim, K.; Zheng, F.; Zeng, Y.; McGregor, M.; Herrington, K. A.; O'Keefe, R.; Jin, N.; VanLandingham, N. K.; Foussard, H.; Von Dollen, J.; Bouhaddou, M.; Jimenez-Morales, D.; Obernier, K.; Kreisberg, J. F.; Kim, M.; Johnson, D. E.; Jura, N.; Grandis, J. R.; Gutkind, J. S.; Ideker, T.; Krogan, N. J., A protein network map of head and neck cancer reveals PIK3CA mutant drug sensitivity. *Science* **2021,** *374* (6563), eabf2911.
4. Zhang, Z.; Rohweder, P. J.; Ongpipattanakul, C.; Basu, K.; Bohn, M. F.; Dugan, E. J.; Steri, V.; Hann, B.; Shokat, K. M.; Craik, C. S., A covalent inhibitor of K-Ras(G12C) induces MHC class I presentation of haptenated peptide neoepitopes targetable by immunotherapy. *Cancer Cell* **2022,** *40* (9), 1060-1069 e7.
5. Takemoto, K.; Matsuda, T.; McDougall, M.; Klaubert, D. H.; Hasegawa, A.; Los, G. V.; Wood, K. V.; Miyawaki, A.; Nagai, T., Chromophore-assisted light inactivation of HaloTag fusion proteins labeled with eosin in living cells. *ACS Chem Biol* **2011,** *6* (5), 401-6.
